# A cluster of three snoRNAs including jouvence required in the gut determines lifespan and confers neuroprotection through metabolic parameters

**DOI:** 10.1101/2025.10.11.681784

**Authors:** Sara Al Issa, Théo Gauvrit, Patricia Daira, Nathalie Bernoud-Hubac, Jean-René Martin

**Affiliations:** Équipe : Imagerie Cérébrale Fonctionnelle et Comportements (ICFC), Institut des Neurosciences Paris-Saclay (Neuro-PSI), UMR-9197, CNRS/Université Paris-Saclay, Campus CEA Saclay, 151 route de la Rotonde, Bâtiment 151 91400, Saclay, France, /; INSA Lyon, CNRS, LaMCoS, UMR5259, 69621 Villeurbanne, France

**Keywords:** snoRNA, aging, longevity, neurodegeneration/neuroprotection, metabolism, triglycerides, cholesterol, gut/brain axis

## Abstract

In our society, the aging of the population is a major concern of public health. Recently we have identified a new snoRNA (jouvence) in Drosophila, and showed that its deletion (F4) reduces lifespan, while its overexpression increases it. F4 deleted flies also present neurodegenerative lesions and a deregulation of metabolic parameters as triglycerides and sterol. However, a deeper characterization of this genomic locus has revealed the presence of two other snoRNAs. Here, we have characterized at the whole organismal level, the role of each them. First, we show that each snoRNAs are expressed in the epithelium of the gut (enterocytes), and in the fat body. Second, in F4 deletion, the re-expression of each snoRNA in the enterocytes or in the fat body is sufficient to improve lifespan, and protect against neurodegeneration in old flies. In addition, depending of the snoRNAs, it rescues the expression of specific deregulated genes within the epithelium of the gut, involved in lipids and sterol metabolism. Consequently, these two metabolic parameters are also rescued, establishing a relationship between the lesions of the brain, the metabolic disorders, the lifespan, and each snoRNAs respectively. Finally, histological stainings as Nile Red and BODIPY C11-581/591 have revealed that the neurodegenerative lesions are due to an increase of free sterol within the brain, and lipid peroxydation in the pericerebral fat body. Altogether, these results point-out a causal relationship between the epithelium of the gut and the neurodegenerative lesions through the metabolic parameters, indicating a gut-brain axis.

## Introduction

In our society, ageing, longevity, and metabolic disorders are major concerns of public health. They result of complex biological processes of accumulation of damages at molecular, cellular, tissues, organs, and whole organism levels (Singh et al., 2019; Fontana et al., 2010). This high complexity related to the fact that neurodegenerative diseases are progressive disorders with typically late-onset increases the difficulty to precisely decipher the genetic and molecular origins of their causes, in other words, where and when they occurred. Genetic, molecular, and physiological approaches have revealed several genes and signaling pathways involved in the determination of ageing, lifespan, and neurodegeneration (López-Otín et al., 2013; 2016; 2023; Gems & Partridge, 2013; Giannakou & Partridge; 2007; Broughton et al., 2005; Owusu-Ansah & Perrimon, 2014; Biglou et al., 2021). Up to now, these mutiple causes have been grouped in 12 hallmarks of ageing (López-Otín et al., 2013; 2023), which reflects their diversity. Among them, the insulin signaling pathway has been the most studied (Giannakou & Partridge; 2007; Broughton et al., 2005; Biglou et al., 2021). Moreover, several studies have underlined the role of lipid metabolism (Johnson & Stolzing, 2019; Liu et al., 2015).

In addition, other metabolic parameters, such as sterol (including cholesterol) homeostasis have also been involved in ageing and neurodegeneration, both in Drosophila (Jing & Behmer, 2020; Tschäpe et al., 2002), as well as in mammals (Saher, 2023). For example, the NPC1b gene promotes sterol absorption from the midgut epithelium (Voght et al., 2007), while the NPC1a causes cholesterol aggregation and age-progressive neurodegeneration (Phillips, et al., 2008). The NPC type C-2 genes family control sterol homeostasis and steroid biosynthesis, and has been used as a model of human neurodegenerative diseases (Huang et al., 2007). To remind, Drosophila cannot synthesize sterols, and thus are dependent of the various forms of sterol contained in the diet (Carvalho et al., 2010). More recently, the microRNAs have also been shown to be major contributors in ageing and neurodegeneration determination (Eacker et al., 2009; Liu et al., 2012; Kato & Slack, 2013; Abe & Bonini, 2013; Soulé et al., 2020; Soulé & Martin, 2020).

Here, we have characterized the role of a new cluster of three snoRNAs including jouvence, in the control of longevity, neurodegeneration and their relationship to lipids and sterol metabolism. First, we show that each snoRNAs is expressed in the epithelium of the gut, and to a less extent in abdominal and pericerebral fat body. Second, in the F4-deleted flies, the re-expression of each of them individually or by group of two or three, either in the epithelium of the gut or in the fat body is sufficient to increase, at various degrees, the lifespan, and to protect against neurodegeneration. At the physiological level, they also differently rescue the metabolic parameters, as triglycerides and sterol. These rescued metabolic parameters are also supported by the restored expression of some specific genes, previously shown to be deregulated (Soulé et al., 2020) within the epithelium of the gut, that are involved in lipids and/or sterol metabolism and associated with each snoRNA. Finally, brain histological stainings, as Nile Red and BODIPY C11-581/591, point-out that the neurodegenerative lesions are due, at least in part, to complex perturbation of sterol homeostasis within the brain, and lipid peroxidation in the pericerebral fat body. Since the snoRNAs are not expressed within the brain, but are required and sufficient in the epithelium of the gut (and too a less extent within the fat body) to prevent the development of neurodegenerative lesions, altogether, these results highlight a causal relationship between the epithelium of the gut and the neurodegenerative lesions linked by metabolic parameters, indicating a gut-brain axis.

## Results

### The [PGal4]4C locus contains a small cluster of 3 snoRNAs

Recently, we have identified a new small nucleolar RNA (snoRNA) (jouvence) in Drosophila, and showed that its deletion reduces lifespan, while its overexpression increases it (Soulé et al., 2020). To achieve this function, jouvence is required and sufficient in the epithelium of the gut. However more recently, a developmental transcriptomic analysis has annotated two additional snoRNAs (snoRNA:2R:9445205, shortly named sno2, and snoRNA:2R:9445410 shortly named sno3) (Graveley et al., 2011) localized just upstream to the first snoRNA:Ψ28S-1153 (jouvence or sno1). Molecular genetic confirms the presence of these two snoRNAs, revealing rather a small cluster of three snoRNAs (Figure 1a). These two snoRNAs, which present a high degree of homology (Soulé et al., 2020) are in accordance to the canonical definition of the H/ACA box snoRNAs (Kiss, 2002; Ye, 2007) since they form a hairpin-hinge-hairpin-tail structure. These three closely located snoRNAs separeted by about only 50 bp, and all included in the F4-deletion, forced us to determine the role of each of them, as well as their putative functional relationship. Using a holistic approach, at the whole organismal level, first we show that each of them are expressed in the enterocytes of the epithelium of the gut, as well as in the ovary, and to a lesser extent in the fat body (Figure 1b). However, their expression levels and patterns slightly differ. As previously shown, jouvence is expressed in all enterocytes of the midgut (Soulé et al., 2020), while sno2 and sno3 are also expressed in the midgut, but not in all enterocytes (Figure 1b). The sno2 and sno3 are also expressed in some cells of the abdominal and pericerebral fat body, while jouvence is only faintly, and not conclusively detectable, by ISH in this tissue. RT-qPCR confirms the expression of each of them, as well as their relative level of expression, both in enterocytes (Figure 1c,d), and in the fat body (Figure 1e). In brief, in the gut, sno2 and sno3 are more expressed than jou, both in young and old flies, as well as in the fat body.

**Figure 1.**
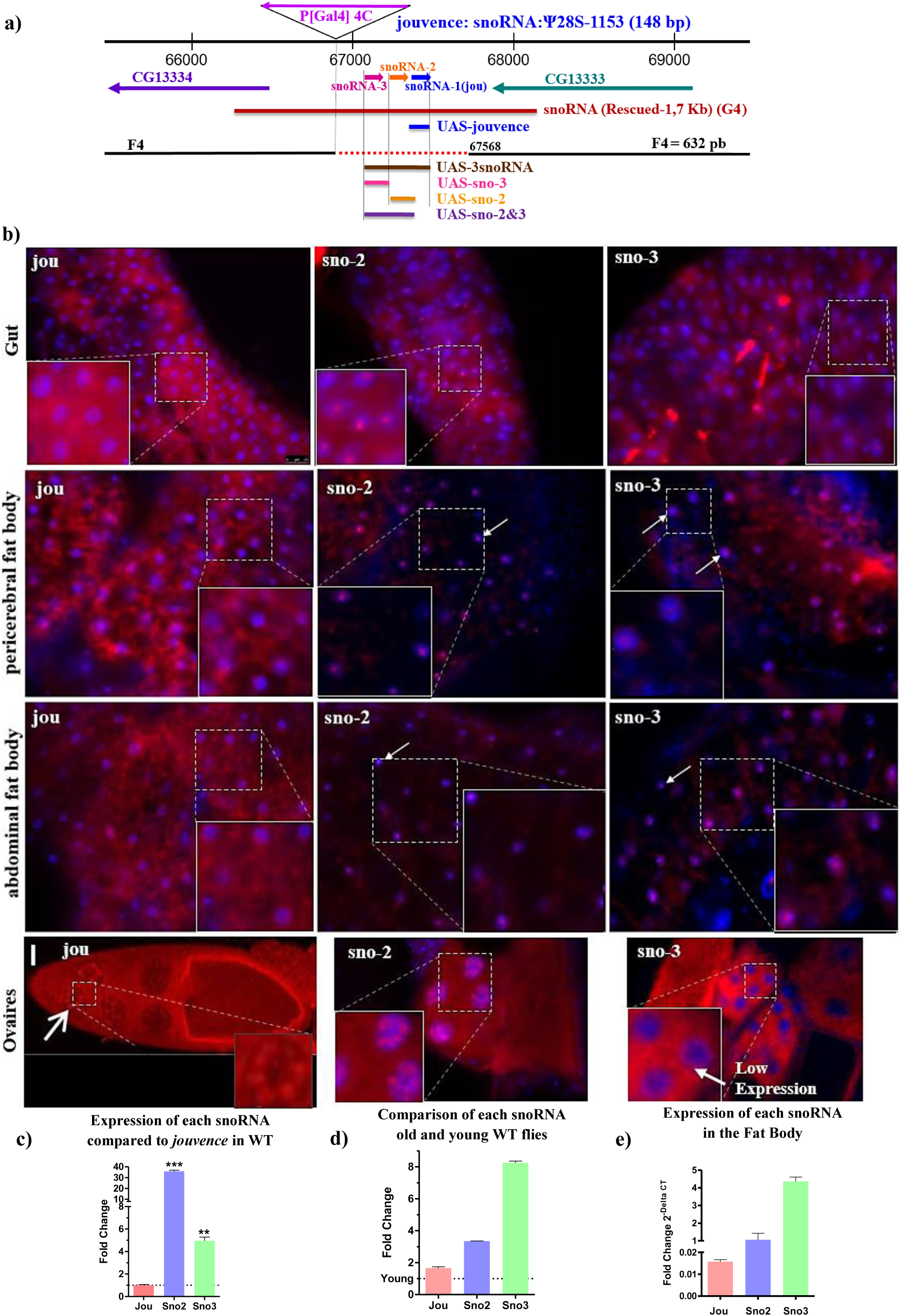
Molecular map and the expression pattern of the 3 snoRNAs. **a)** The snoRNA:Ψ28S-1153 (jouvence or sno-1) and the two other snoRNAs (sno-2 and sno-3) are in the inverse orientation of the two encoding genes: CG13333 and CG13334. The deletion (F4) of 632 bp (red dotted line) encompasses the 3 snoRNAs. The 1723 bp genomic DNA fragment (red bar) (G4) used to generate transgenic flies. The snoRNA fragment (blue bar) of 148bp used to generate UAS-jou construct, the 166 bp fragment (orange bar) corresponding to UAS-sno-2, the 157 bp fragment (pink bar) corresponding to UAS-sno-3, the fragment (purple bar) corresponding to UAS-sno-2&3, and the brown bar fragment to UAS-3snoRNA. **b)** *In situ* Hybridization (ISH) of each snoRNA (jou, sno-2, sno-3) on whole fly (Cryostat section) reveals a restricted expression in the epithelium of the gut, in the pericerebral and the abdominal fat body (snoRNA = red dots = white arrow). Each of them are also expressed in the nurse cells of the ovary (for jouvence, see also Soulé et al., 2020). Blue corresponds to DAPI staining (scale bar = 25μm). Inset (magnification) showing that each snoRNA are located in the nucleolus. **c,d,e)** RT-qPCR (Taqman) showing the expression level of each snoRNAs on dissected gut of Wild-Type CS flies. **c)** In young (7 day-old), the snoRNA levels are normalized to the level of rp49. The expression level of jou was used as a reference and set to 1. **d)** In 40 day-old flies, the snoRNA levels are normalized to the level of rp49 and are presented relative to the levels observed in 7-day-old flies, used as the reference (set to 1, represented by dotted line on the histogram). **e)** in fat body, RT-qPCR (Taqman) showing the expression level (fold change - delta CT, of each snoRNAs on dissected abdominal fat body. Three independent biological replicates were done (n=3). Statistics: compared to jouvence level used as reference = 1. (p-values) (* p<0,05; ** p<0,005; *** p<0,0005). Errors bars represent the mean +/- S.E.M. (p-value were calculated using the student’T test, using Prism).

### Targeted expression of each snoRNA increases lifespan

In order to determine which of the snoRNAs, or two or three of them, are required to rescue longevity, we use three P[Gal4] drivers; Myo1A-Gal4 and two inducible Gene-Switch (Mex-GS and CG89978-GS) (see Suppl. Figure 1 for their expression pattern) for which the activity can be induced only in adulthood by feeding the fly with Mifepristone (RU486). With Myo1A-Gal4, specifically expressed in enterocytes (Jiang et al., 2009), the expression of each snoRNA alone, as jou, sno2 or sno3, increases the longevity (Figure 2a). The expression of two (sno2 and sno3), or three (jou, sno2 and sno3) also similarly increases the longevity, indicating that each snoRNA itself is sufficient to increase lifespan. Since Myo1A-Gal4 is also transiently expressed during development (Morgan et al., 1994), to overcome the possibility that these effects could be due to a developmental effect, we used two inducible drivers, as Mex-GS and CG8997-GS. Mex-GS (Soulé et al., 2020) is expressed in several enterocytes, but not exclusively, as suggested by a transcriptomic analysis performed on the epithelium of the gut Dutta et al., 2015). Nevertheless, the expression of each snoRNA individually, only in adulthood, increases lifespan, although the rescue is not as important as with the Myo1A for jou (Figure 2b-g). The RU486 itself does not have any effect on lifespan (Figure 2b), while the expression of the sno2&3 (Figure 2f), or the 3snoRNAs (Figure 2g) yields a slightly different effect compared to Myo1A (Figure 2a), but nevertheless, still increases lifespan. Similarly, using the CG8997-GS, expressed mainly in the R3 region of the gut, also leads to an increase of lifespan (Suppl. Figure 2a-e).

**Figure 2.**
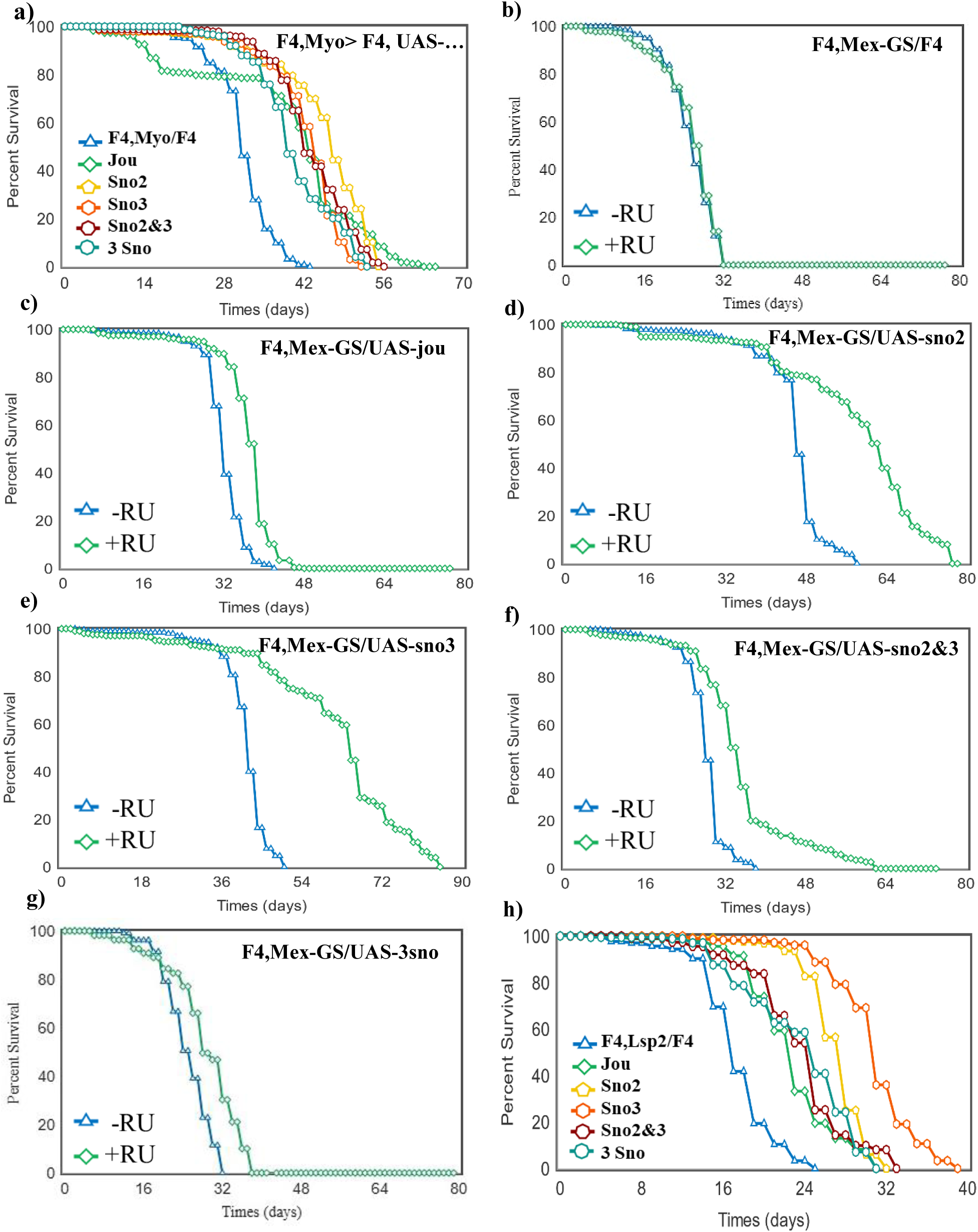
Targetted expression of each snoRNA in the enterocytes or in the fat body increases lifespan. Longevity test results (survival curve - decreasing cumulative) of the targeted expression of each snoRNA specifically in the enterocytes or in the fat body in F4 flies compared to the F4 flies whose don’t express any snoRNA. **a)** Myo1A-Gal4 driving each UAS-snoRNA in the enterocytes. **b)** F4,Mex-GS/F4 fed with RU486. **c,d,e,f,g)** Mex-GS driving each UAS-snoRNA in enterocytes fed with RU486. **h)** Lsp2-Gal4 driving each UAS-snoRNA in the abdominal and pericerebral fat body. For the number of flies, age in days at % mortality, and detailed statistics, see Supplementary Table-S2. p-value calculated by log-rank test using OASIS Sofware.

Since ISH, complemented by RT-qPCR (Figure 1), reveal that the snoRNAs are also expressed in the fat body, the re-expression of each of them using the Lsp-Gal4 driver increases lifespan (Figure 2h) to various degree, as well as the expression of two or three of them. Interestingly, the re-expression of sno3 is by far the most efficient. Finally, since the Lsp-Gal4 is also known to be expressed during development (Benes et al., 1996), we used the Lsp-GS to express the snoRNA only in adulthood to circumvent any putative developmental effect. The expression of jou importantly increases lifespan, sno2 has a moderate effect, sno3 does not have any effect (even has a tendency toward a negative effect), while the sno2&3 together presents a moderate effect similar to sno2 alone (Suppl. Figure 2f-j). In regard to the effect obtained with Lsp-Gal4 driving sno3 (Figure 2h), these results suggest that the sno3 does not have any effect in the fat body in adulthood, and that only jouvence and the sno2 are active in the fat body in adult. It also argues that the previous effect observed with the sno3 (Figure 2h) is likely due to a developmental effect.

### Targeted expression of each snoRNA prevents neurodegenerative lesions

Since the re-expression of each snoRNA in the F4-deleted flies increases lifespan, we wonder if in parallel, the neurodegenerative lesions observed in old flies are also rescued or at least improved, indicating a putative neuroprotection. As previously reported (Soulé & Martin, 2020), the deletion of the 3 snoRNAs (F4) increases the number of neurodegenerative lesions (vacuoles) within the brain in old flies (Figure 3). Myo-Gal4 driving the expression of jou within the enterocytes rescues the number of lesions (Figure 3a), as well as the total surface area of these lesions (Suppl. Figure 3a). We observe similar effects with the sno2, sno3, sno2&3, and even a stronger effect with the 3snoRNAs (Figure 3a). However, with the sno2, sno3 or sno2&3, the surface area are not rescued, while they are well rescued with the 3snoRNAs (Suppl. Figure 3a). These results suggest that jou is the most efficient to rescue the neurodegenerative lesions, while when accompanied by sno2 and sno3 (3snoRNAs altogether), the rescued effect is even better (both for the number of vacuoles and the surface area). Similar results are observed when each snoRNA are expressed only in adulthood, with Mex-GS (Figure 3b) and CG8997-GS (Figure 3c), again here, with the stronger effect with the 3snoRNAs, although a slight improved effect is also observed without RU486 induction (see below for further explanation). For the surface area (Suppl. Figure 3b,c), similar results are obtained with jou, while slight improvements are observed with sno2, sno3, sno2&3 or the 3snoRNAs (again here, see below for further explanation). Altogether, these results obtained with three different drivers indicate, although with some differences, that the expression of the snoRNAs within the enterocytes is sufficient to rescue the neurodegenerative lesions in old flies. Since the snoRNAs are also expressed in the fat body, we use the Lsp-Gal4 line. All of them improve (partially rescue) the number of lesions (Figure 3d), and the surface area (Suppl. Figure 3d). Again here, the strongest effect is observed with the expression of the 3snoRNAs. Thus, the expression of each snoRNA within the fat body also prevents the neurodegenerative lesions, suggesting that the gut defect could be compensated by a rescue in the fat body.

**Figure 3.**
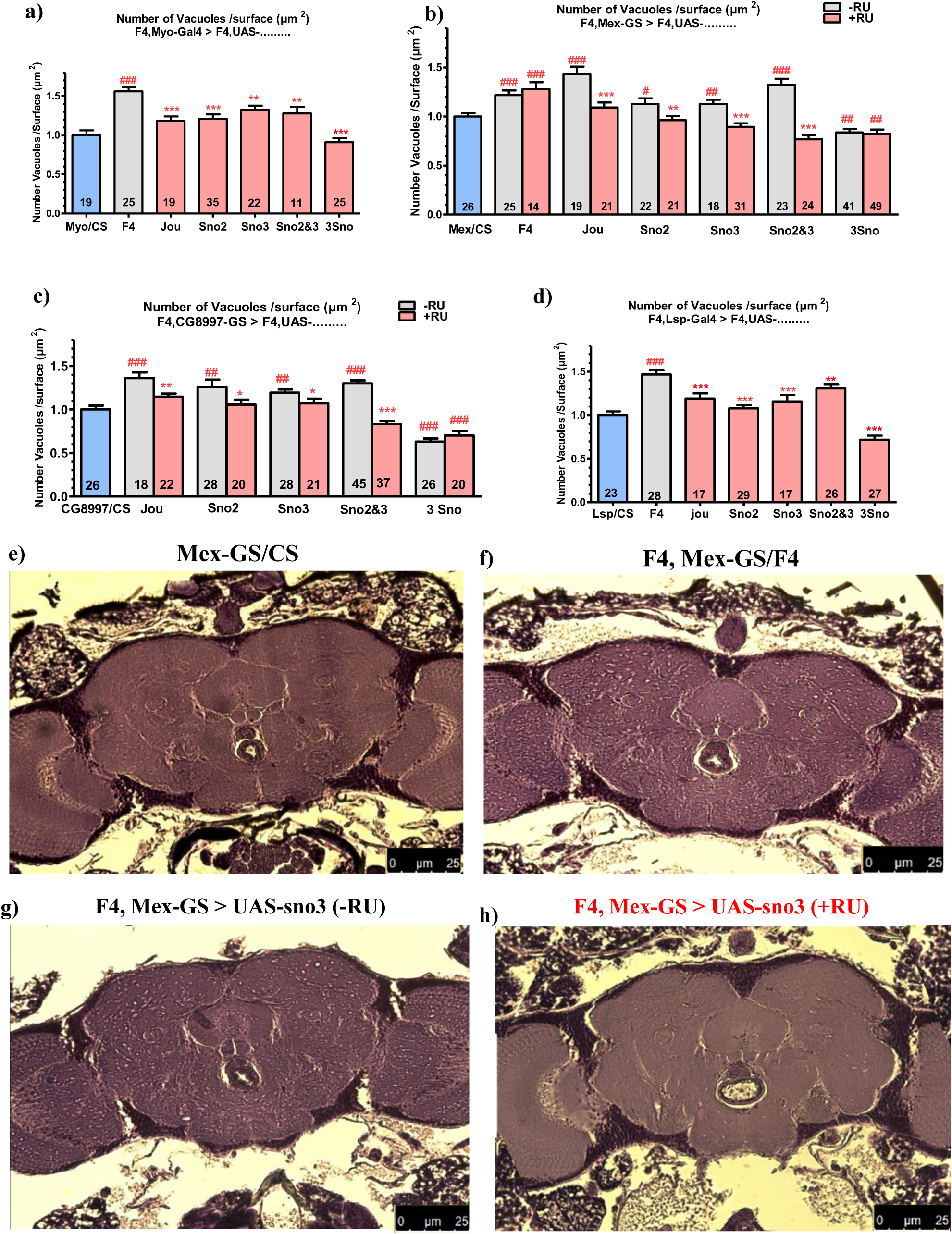
Quantification of vacuoles of neurodegenerative lesions in old flies. Each expression (rescue) is performed in the F4-deleted genetic background. **a)** Number of vacuoles for the Myo1A-Gal4 driving each UAS-snoRNA in the enterocytes. **b)** Number of vacuoles for the Mex-GS driving each UAS-snoRNA in the enterocytes of F4-deleted flies, without and with feeding RU486 to induce the expression of the UAS-snoRNA transgenes. **c)** similar, but using the driver line CG8997-GS. **d)** similar, but using the driver line Lsp2-Gal4 to target the snoRNA in the fat body. Numbers within the histograms indicate the number of flies, errors bars represent the mean ± S.E.M. p-value were calculated using the student-T test (Prism). Asterisks indicate significant differences compared to flies that never received RU (gray), while hashtags (#) highlight significant differences compared to wild-type flies (Blue) (Myo/CS, Mex/CS, CG8997-GS/CS, Lsp/CS) (*p<0.05; **p<0.005 ; ***p<0.0005). **e,f,g,h)** Micro-photography showing the vacuoles in old flies of a Wild-Type fly (e), in a F4-deleted fly (f), in a F4,Mex-GS>UAS-sno3, without RU (g), and after induction with RU (h).

### RT-qPCR confirms the expression of each snoRNA by P[Gal4] lines

To validate that the snoRNAs are well expressed in the gut or in the fat body, we have performed RT-qPCR. With Myo1A-Gal4, jou is overexpressed by a factor of 500, while it is of 100 with the 3snoRNAs (Figure 4a). For the sno2, the overexpression is only of 30 fold, while it is of 350 fold with the sno2&3, and 400 fold with the 3snoRNAs (Figure 4b). For the sno3, the overexpression is exactly of one (perfect rescue level), while it is of 7 fold with the sno2&3, and 6 fold with the 3snoRNAs (Figure 4c). Interestingly, although the UAS-construct is the same for each snoRNA, and the transgenic insertion site within the genome is the same for the sno2, sno3, and sno2&3 (VK27, 89E11, chromosome 3L), thus, *per se*, excluding an insertion site effect, we observe a strong difference in the expression level of the snoRNAs under the control of Myo1A-Gal4. Thus, this difference is unlikely due to a difference in the transcription level, but likely rather to the stability of the snoRNA itself, a question that remains to be determined. Nevertheless, the presence of the sno3 (Figure 4c) seems to be the most important to precisely rescue, almost like in WT flies, its expression level.

**Figure 4.**
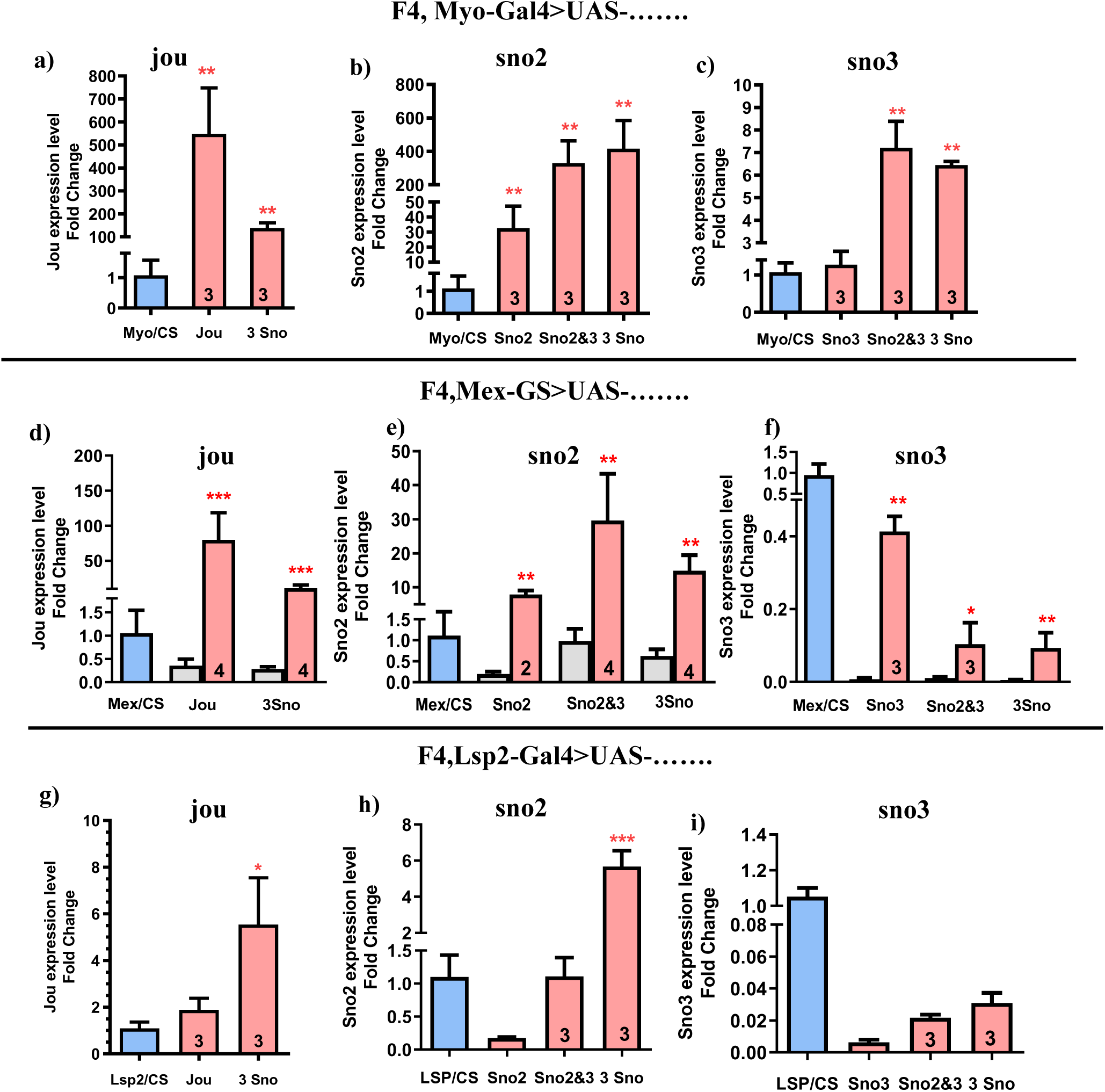
Expression level of each snoRNAs (RT-qPCR) in rescued F4 flies. **a,b,c)** RT-qPCR (Taqman) results of jou (a), sno2 (b), or sno3 (c), following the expression of the appropriated snoRNA by the Myo1A-Gal4 driver in the enterocytes (for example, in a, the quantification of the sno2 and sno3 have not been performed since the flies are in F4 deleted genetic background, and thus they are absent, idem of the other genetic combinations). **d,e,f)** idem for the Mex-GS driver in the enterocytes, with and without RU486. **g,h,i)** idem for the Lsp2-Gal4 driver in the abdominal and pericerebral fat body. Statistics: for each snoRNA, comparison to control Myo/CS, Mex/CS and Lsp/CS used as reference (=1). (p-values) (* p<0,05; ** p<0,005; *** p<0,0005). Errors bars represent the mean +/- S.E.M. (p-value were calculated using the student T test, using Prism).

With the inducible Mex-GS, we observe quite similar pictures, but with a general lower expression level compared to Myo1A-Gal4 (Figure 4d-e-f). For example, the level of jouvence is 100 fold compared to 400 for the Myo1A-Gal4, and so on for the sno2 (Figure 4e) and sno3 (Figure 4f). However, the RT-qPCR quantification has revealed that in all the Mex-GS>UAS-snoRNA (any of them), we detect more or less, a slight level of the snoRNA (Figure 4d-e-f, in grey), without feeding the flies with RU486, suggesting a weak leakage of the snoRNA. If this leakage is due to the Gene-Switch that is not well locked – a phenomenon already reported for several Gene-Switch lines (Poirier et al., 2008), or due to the leakage of the UAS-snoRNA itself, remains to be precisely determined. Whatever the case may be, all of these variabilities in the overexpression level of the snoRNAs, and/or the leakage of them, likely contribute to the difference observed in the degree of rescue of longevity and/or neurodegenerative lesions.

Concerning Lsp2-Gal4, the level of jou is doubled, compared to control flies (Figure 4g), while is it increased by 6 fold for the 3snoRNA. For the sno2, the level is only 0,1 compared to control (Figure 4h), while it is one for the sno2&3, and 6 fold for the 3snoRNAs. For the sno3, as for the sno2, it is only 0,1 compared to control (Figure 4i), while it is 0,2 for the sno2&3 and 0,3 for the 3snoRNA. Here, although this quantification has been done on whole flies, obviously the expression level is much lower than with the other gut drivers. Moreover, the expression level is clearly very different for jou, sno2 and sno3, which again could be responsible for the difference observed in the phenotype (longevity and brain lesions).

### Targeted expression of each snoRNA rescues triglycerides and sterol

To characterize the neurodegenerative lesions within the brain, and notably to check if these lesions could be due to apoptosis, we have performed immuno-histological stainings with an anti-caspase-3 antibody, which reveals several aggregates and a hypertrophy of the pericerebral fat body (Figure 5a) in F4-deleted flies, without any striking staining within the brain. To investigate further, we have quantified, in the whole fly, the triglycerides (TG). First, in F4-deleted flies with Myo1A-Gal4, the level of TG is increased (Figure 5b). The expression of jou in enterocytes rescues the level of TG. Similarly, the expression of sno2, or sno3, or sno2&3, or the 3snoRNAs leads to similar results, with a maximum effect observed with the sno3. Similar effects were obtained when the snoRNAs are expressed only in adulthood using Mex-GS after RU486 feeding (Figure 5c). The expression of each snoRNA in the fat body using the Lsp2-Gal4 (Figure 5d) also rescues (decreases) the level of TG, again, with a maximum effect observed with the sno3.

**Figure 5.**
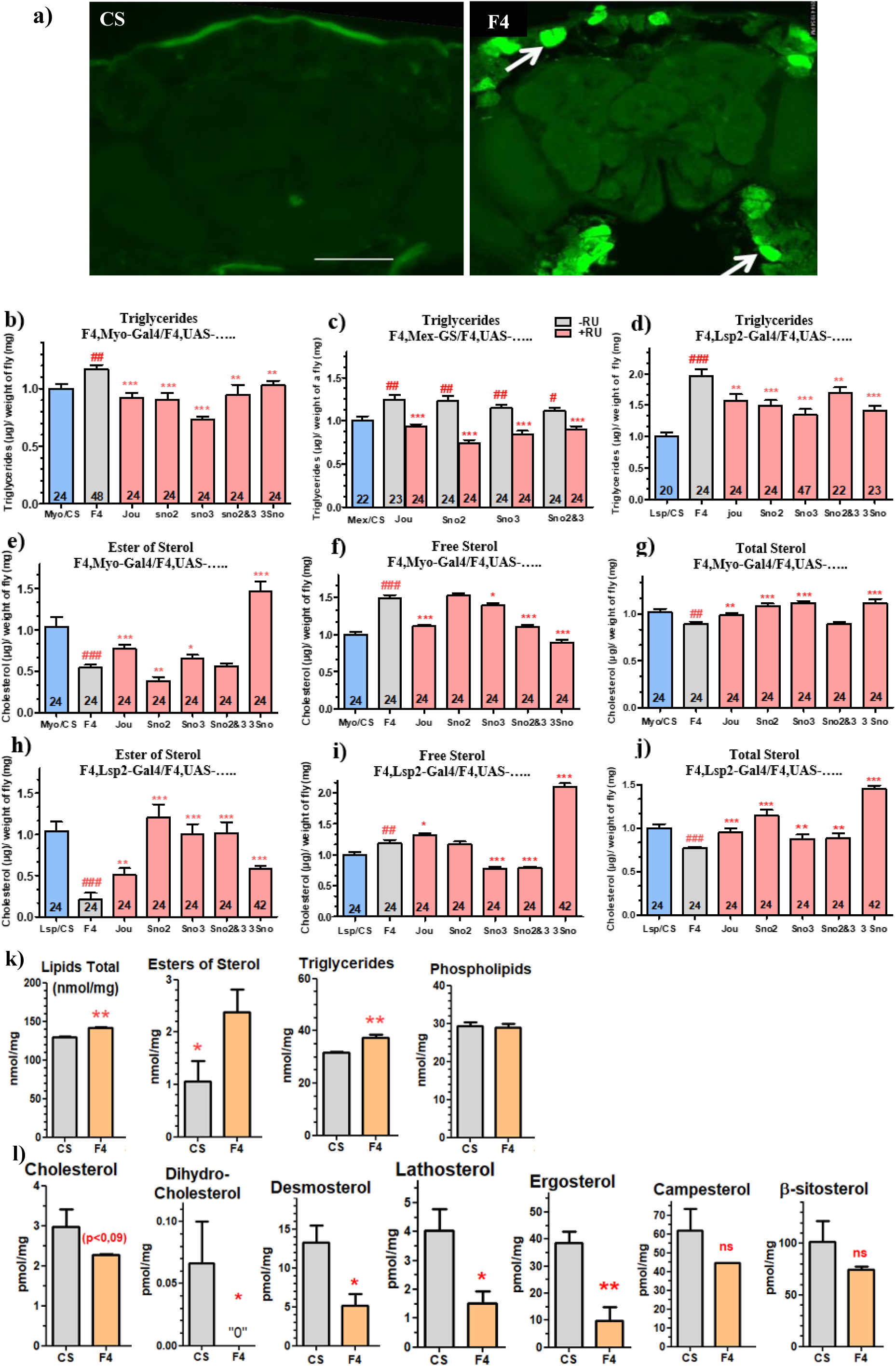
Deregulated Triglycerides and Sterols are rescued by targetted expression of snoRNA. **a)** Immuno-Histological stainings using anti-caspase-3 antibody on 40 day-old flies, reveal aggretates located in the pericerebral fat body (F4-deleted flies) compared to Wild-Type (CS) controls flies (scale bar = 100µm). **b)** TG dosage on whole flies (7 day-old), following the expression of each snoRNA using Myo1A-Gal4 line. **c)** Dosage of TG using Mex-GS, with and without feeding RU486. **d)** Dosage of TG following the expression of each snoRNA in the abdominal and pericerebral fat body, using Lsp2-Gal4. Quantification of sterol-ester **(e)**, free **(f)**, and total **(g)** levels in wild-type controls (CS), F4-deleted flies (F4), and flies expressing each snoRNA in the enterocytes of the gut under the control of Myo1A-Gal4. Quantification of sterol-ester **(h)**, free **(i)**, and total **(j)** levels in flies expressing each snoRNA in the fat body under the control of Lsp2-Gal4. Numbers in the histograms indicate the number of flies, errors bars represent the mean ± S.E.M (*p<0.05 ; **p<0.005 ; ***p<0.0005). P-value were calculated using the student T test (Prism). Asterisks indicate significant differences compared to flies that never received RU (gray), while hashtags (#) highlight significant differences compared to wild-type flies (Blue) (Myo/CS, Mex/CS, CG8997-GS/CS, Lsp/CS). **k)** Lipidomics analysis reveals the precise amount of each class of lipids. **l)** Amounts of different classes of sterol. All of them are decreased, although the campesterol and β-sisterol are not statistically different. Remarks that the Y-scales are different for each sterol, and notably that the phytosterol (ergosterol, campesterol and β-sisterol) are much higher.

To go further, we wondered if the sterols, another major components of the metabolism, were also deregulated. We have quantified the sterols on the whole flies, as for the TG. The ester of sterol is strikingly reduced in F4-deleted flies compared to control (Figure 5e), the free sterol is increased (Figure 5f), while the sum of both, the total-sterol, is just slightly decreased (Figure 5g). The expression of jou in the enterocytes (Myo1A-Gal4) rescues the sterol-ester and the free sterol (Figure 5e), while the expression of the sno2 don’t have any effect on sterol (Figure 5f). The expression of sno3 has just a mild rescue effect (Figure 5e,f,g), whereas the expression of sno2&3 don’t have any effect on sterol-ester, but solely on free sterol (Figure 5g). Finally, the expression of the 3snoRNAs rescues the sterol-ester and the free-sterol (Figure 5e,f). In the fat body, the expression of sno2, sno3 or sno2&3 rescues the sterol-ester and the free sterol except sno2 (Figure 5h,i). For jou and the 3snoRNAs, the ester of sterol is partially rescue with any rescue effet on free sterol (Figure 5h,i). In conclusion, strikingly, the main effects on sterol (both -ester and -free) are due to the presence of jou in the gut, either when it is expressed alone or when it is included in the cluster of the 3snoRNAs. However, in the fat body, the primary effect on sterol (both esterified and free) results of sno3, either alone or co-expressed with sno2.

### A lipidomic analysis reveals different classes of lipids and sterols

To investigate further the nature of the deregulated lipids and sterols, as well as their fatty-acid components, we performed a lipidomic analysis. In F4-deleted flies, we observe an increase in total lipids, triglycerides, and ester of sterol, while the amount of phospholipids is not modified (Figure 5k). Inversely, the different forms of sterol, as the cholesterol, dihydro-cholesterol, desmosterol, lathosterol, and ergosterol are importantly decreased, while the campesterol and the β-sitosterol are also decreased, but these last are not statistically differents (Figure 5l) (remark that the levels/quantity of phystosterols as ergosterol, campesterol and β-sitosterol are much higher that the others).

In contrast, the precise determination of the fatty-acid (FA) components is more complex. In overall, for the total lipids and triglycerides, the short FA, as 12:0 and 14:0 are increased, while the longer ones (>16:0) are decreased, with some exceptions (Suppl. Figures 4&5). For the sterol-ester, it is quite similar, except that the 16:0 are increased (suppl. Figure 6). Finally, for phospholipids, roughly, there is no striking modification except a decrease in 14:1 (Suppl. Figure 7). Altogether, these results indicate that the deletion of the 3 snoRNAs (F4) perturbs quantitatively some specific FA, and more especially the short ones (12:0 and 14:0).

### Neurodegenerative lesions are due to peroxidized lipids and sterol deregulation

To gain insigth into the nature of the brain lesions and in accordance with the deregulated general metabolic parameters (TG and sterols), we performed different histological stainings on the brain. Lipid stainings using Nile Red reveal several positive puncta (or aggregates) spreaded all over the brain in F4-deleted flies compared to control (Figure 6a-b), reminding the vacuoles observed in the parrafin section of the old flies, ascribed to neurodegenerative lesions (Figure 3). To determine if these points/aggregates are associated to glial cells or to neuronal cells, similar stainings were performed using repo-Gal4>UAS-GFP to label the glial cells, and the n-syb-Gal4>UAS-GFP to label the neuronal cells. Double-stainings reveals that the Nile-Red overlaps with the neuronal tissue but not with the glial cells (Suppl. Figure 8). Since Nile-Red is a fluorescent lipophilic dye characterized by a shift of emission from red to yellow according to the degree of hydrophobicity of lipids [34], we took advantages of this unique property to quantify the ratio of red and yellow emission in order to discriminate the labeled lipids. According to a lipids reference (Diaz et al., 2008), the Nile-Red aggregates in F4-deleted flies correspond to an increase of free sterol to the detriment of sterol-ester (Figure 6a-c).

**Figure 6.**
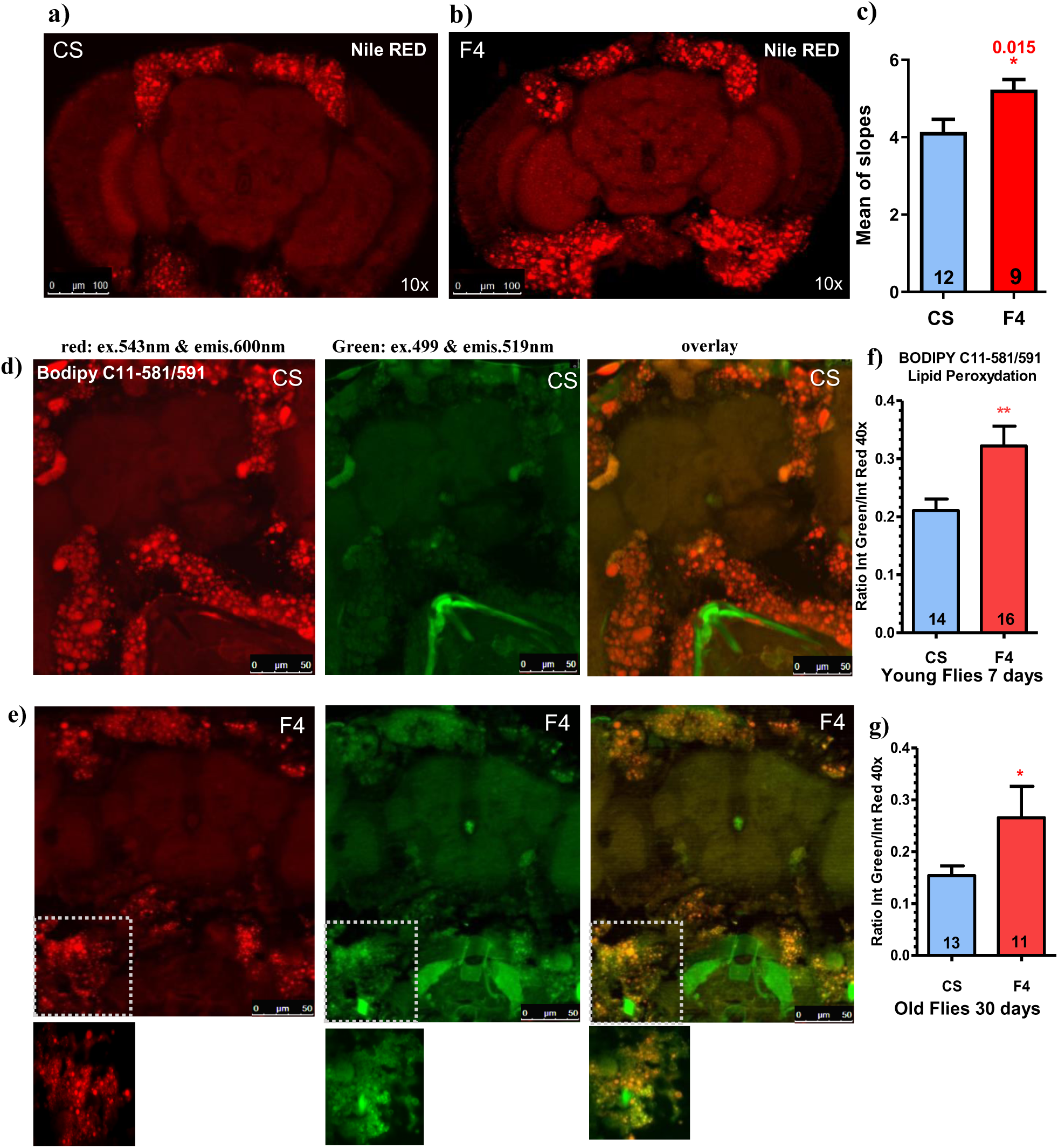
Nile Red and BODIPY C11-581/591 highlight the nature of the neurodegenerative lesions. **a,b)** Nile Red staining of the brain shows the pericerebral fat body in Wild-Type CS (a) and F4-deleted flies (b). Several unique puncta (or aggregates) are visible, principally in F4 flies (b). The double nature of the Nile Red, a shift of emission from red to yellow according to the degree of hydrophobicity of lipids, allows to discriminate the nature of the lipids. In **(c),** the mean of the slope indicates a higher amount of free-cholesterol in the aggregates. **d,e,f,g)** BODIPY C11-581/591 staining of the brain shows the pericerebral fat body in Wild-Type CS (d) and F4-deleted flies (e). First column, in red: excitation at 543 nm & emission at 600-650 nm. Second column, green: excitation at 499 nm & emission at 519 nm. Third column, overlay. The quantification of the ration green/red confirms that F4 flies have more peroxidized lipids than control flies, both in young **(f)** and old flies **(g).**

As lipids could be peroxidized, we also measured the lipid peroxidation using the BODYPY C11-581/591, which reveals the non-peroxidized lipids in red, while the peroxidized ones are in green (Bailey et al., 2015). Figure 6d shows that the control have just few green staining, whereas F4 (Figure 6e) have more intense green staining. The quantification of the ratio green/red reveals that the F4-deleted flies have more peroxidized lipids that control, both in young and old flies (Figure 6f-g) specifically in the pericerebral fat body.

### Targeted expression of each snoRNA within enterocytes differently rescues the expression level of genes involved in lipid metabolism

To temptingly decipher the molecular mechanisms linking the epithelium of the gut, the metabolic parameters, and the neurodegenerative lesions, we quantify by RT-qPCR some selected genes involved in the metabolism of lipids and sterols. Based on the previous transcriptomic analysis performed on the dissected gut (Soulé et al., 2020), first, we have analysed NPC1, NPC2, and ninaD, three genes involved in the sterol (cholesterol) metabolism. NPC1, decreased in F4, is partly rescued by the expression of sno2, sno3 and sno2&3, but not by the 3snoRNAs neither by jou (Figure 7a). NPC2, also decreased in F4, is well rescued by the re-expression of sno2, sno3, sno2&3, and the 3snoRNAs, but not by jou (Figure 7b). ninaD, which inversely, the expression is increased in F4 (Figure 7c) (Soulé et al., 2020; Soulé & Martin, 2020), only the re-expression of the sno3, or sno2&3, or the 3snoRNAs rescues its mRNA level. Altogether, these results indicate that the rescue of the mRNA level of NPC1 or NPC2 depend of the sno2 and/or the sno3, but not jou, while ninaD is achieved solely by the sno3, since the presence of the sno3 is mandatory to rescue it (sno3, sno2&3, 3snoRNAs).

**Figure 7.**
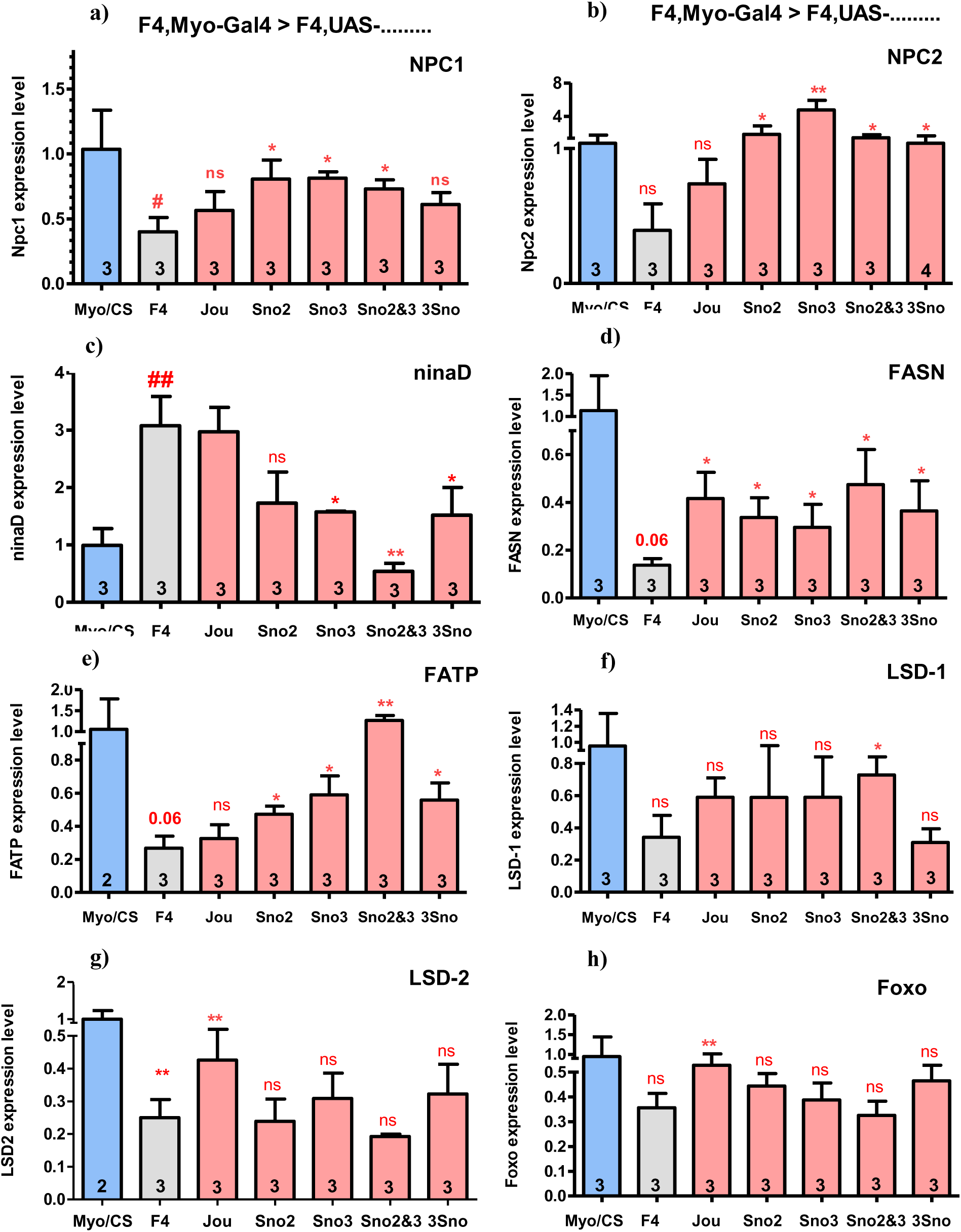
Different genes are differentially rescued by the snoRNA (RT-qPCR). RT-qPCR (SybGreen) of 8 different encoding genes performed on dissected gut of Controls, F4-deleted flies and targeted expression of each snoRNA in the enterocytes driven by Myo1A-Gal4. **a)** NPC1. **b)** NPC2. **c)** ninaD. **d)** FASN. **e)** FATP. **f)** Lsd1 (perilipin-1). **g)** Lsd2 (perilipin-2). **h)** FOXO. Statistics: The expression level of each gene in Control Wild-Type (Myo1A/CS) is used as reference (fixed to 1). Three independent biological replicates (n=3). (p-values) (* p<0,05; ** p<0,005; *** p<0,0005). Errors bars represent the mean +/- S.E.M. (p-value were calculated using the student T test, using Prism).

We also investigate some genes involved in the fatty-acid and triglycerides metabolism. The gene FASN (fatty-acid synthase), decreased in F4 (Soulé et al., 2020), is partly rescued by the re-expression of any snoRNA (jou, sno2, sno3, sno2&3, and 3snoRNAs) (Figure 7d). FATP (fatty-acid transporter protein), also decreased in F4, is not rescued by jou, but is partly rescued by the re-expression of the sno2, sno3, and the 3snoRNAs, while it is perfectly rescued by the expression of the sno2&3 (Figure 7e). We also check the expression level of the two perilipin genes (Lsd1 and Lsd2), which are known to be crucial for the formation of lipids droplets (Beller et al., 2010; Bi et al., 2012). While both are decreased in F4 (Figure 7f-g), only Lsd1 is rescued by the sno2&3, while Lsd2 is rescued only by jou. Strangely, for the two perilipins, the lack of rescue following the re-expression of each snoRNA in the gut seems to be, at least partly, due to a larger variability observed between the three independent replicats (although these are the same RNAs used for the other genes).

We also quantify the gene FOXO, one of the crucial gene of the insulin-signaling pathway (Bolukbasi et al., 2017). FOXO is decreased in F4, confirming the RNA-Seq (Soulé et al., 2020), while it is only rescued by the re-expression of jou (Figure 7h). Interestingly, this rescue-pattern is an inverted mirror image of the rescue-pattern observed for the ninaD gene. Similar results obtained by RT-qPCR on the same genes have been observed with the Mex-GS line after RU486 feeding, indicating that the expression of these genes within the epithelium of the gut only in adulthood is sufficient (Suppl. Figure 9). In complement, the same genes have been investigated using the Lsp2-Gal4 (fat body) (Suppl. Figure 10). In brief, NPC1 and NPC2 are not rescued, while the expression of jouvence nicely rescues the ninaD level, almost as a mirror image of the gut one’s. For the FASN and FATP, the rescue is quite similar to the gut lines (although with some slight differences). Lsd2 is not rescued, as with the gut lines, except with the 3snoRNAs. Finally, for FOXO, the results present some variabilities, due to a large difference between the replicates.

## Discussion

We have reported that the deletion of a cluster of 3 snoRNAs including jouvence (F4-deletion) decreases the longevity, while the re-expression of each of them individually, or two or all 3 together, specifically within the enterocytes rescues it to various degree. These rescues have been demonstrated with three independent driver lines: Myo1A-Gal4, Mex-GS, and CG8997-GS. Although the expression pattern of the two last does not completely cover the entire midgut, as Myo1A-Gal4, nevertheless, the longevity is improved, indicating that the re-expression of the snoRNA only in adulthood is sufficient to rescue the longevity. Moreover, as revealed by the ISH and corroborated by RT-qPCR (Figure 1), the 3 snoRNAs are also expressed in some cells of the abdominal and pericerebral fat body. Astonishingly, the re-expression of each snoRNA individually, or two, or three together, by the Lsp-Gal4, or induced only in the adulthood by the Lsp-GS (Suppl. Figure 2) also improves lifespan.

Since the snoRNAs overexpressing flies live longer, we wonder if these long-lived flies could have more or less brain defects. F4-deleted old flies present more neurodegenerative lesions (vacuoles) than the control (which also present some lesions), as previously reported (Soulé & Martin, 2020). However, the re-expression of each snoRNA (or 2 or 3) within the gut rescues both the number (Figure 3) and the surface are of neurodegenerative lesions (Suppl. Figure 3). As for the longevity, these effects are also obtained with 3 independent gut drivers, which among them, two are expressed (induced) only in adulthood, indicating that the expression of the snoRNA is sufficient only in adulthood. Moreover, the expression of each snoRNA within the fat body (Lsp-Gal4) is also sufficient to improve the neurodegenerative lesions (both the number and the surface area) with a better effect when we express the 3 snoRNAs altogether. Therefore, these results indicate that the defects induced by the missing snoRNA within the enterocytes could be compensated by the re-expression of them in the fat body. They also reveal a correlation (though it is not a perfect linear correlation) between the longevity of the flies and the severity of the brain lesions.

Since none of the snoRNAs are expressed within the brain, to trace a causal relationship between the tissues that express the snoRNA (the gut and to less extent, the fat body) and the brain lesions, some metabolic parameters have been quantified. These experiments were firstly guided by the fact that immuno-histological stainings of the heads and brains have revealed that the main visible histological lesions were localized in the pericerebral fat body, indicating notably a hypertrophy of the fat body (Figure 5). Thus, while the TG are increased in F4, a normal level of TG is rescued to various extents by the different snoRNA when they are expressed alone or two ot three together within the enterocytes (Figure 5). Similar effects (although with some variabilities) are also obtained with the expression of the snoRNAs within the fat body. For the sterols and their various forms, the sterol-ester seems to be the most affected (a decrease) in F4, while the free sterol are increased, for a total sterol being only slightly affected (Figure 5). However, in contrast to the TG, which were rescued by all the snoRNAs, the sterol and notably the sterol-ester are rescued solely by jou (either alone or included within the 3 snoRNAs). Interestingly, these last results also correlate to the longevity and the neurodegenerative lesions. Therefore, they allow to suggest that the metabolic parameters (Tg and sterol) could fulfill the gap between the gut (or the fat body) and the neurodegenerative lesions. In other words, the deregulation of the metabolic parameters due to the missing snoRNA, primarily within the gut and in relay within the fat body, chronically leads to neurodegenerative lesions.

To characterize the nature of the brain lesions, different stainings have been performed. First, the Nile Red, known to stain the neutral lipids (Diaz et al., 2008) has revealed several small aggregates within the neuronal tissues, as confirmed by the double staining with n-Syb-Gal4 (Suppl. Figure 8). Moreover, as the Nile Red fluoresces in different wavelength, from red to yellow in relation to the degree of hydrophobicity of lipids, this last property has suggested that these aggregates are enriched in free sterol. Second, the BODIPY C11-581/591, which fluoresces differentially according to the degree of peroxidation of lipids, has nicely reveals a higher degree of lipid peroxidation in F4, especially in the pericerebral fat body. Altogether, these various stainings, either in the fat body or the brain, conjugates to the deregulation of the general metabolic parameters (TG and sterol) suggests that the neurodegenerative lesions might be a consequence of the chronic deregulation of the metabolic parameters induced by the missing expression of the different snoRNAs, occurring in the gut and/or the fat body.

To strengthen this causality, and notably to decipher the relation between the snoRNA within the enterocytes and the deregulated metabolic parameters, we have quantified the expression of several genes involved in the lipid and sterol metabolism from the dissected gut of various snoRNA rescued genotypes. In agreement to the previous transcriptomic analysis (Soulé et al., 2020), we have confirmed that the re-expression of all snoRNAs, except for jou, is sufficient to increase (partly rescue) the level of the NPC2a gene, one of the main gene regulating cholesterol homeostasis (Huang et al., 2007). This is also similar for the NPC1a, a gene involved in intracelular sterol trafficking (Phillips et al., 2008). However, ninaD is not rescued by jou, but instead it seems to require the presence of the sno3, since it is rescued by sno3 alone, or sno2&3 or by the 3snoRNAs. Altogether, these results observed with three main genes involved in sterol homeostasis leading to chronic sterol metabolic defects likely participate to neurodegeneration. Concerning FASN and FATP, two genes involved in the fatty-acid and triglycerides synthesis and homeostasis (Heier & Kühnlein, 2018), they are also rescued by all the snoRNAs, except for FATP with jou. However, it is not the same for all analysed genes, as for example, the two perilipins (Lsd1 and Lsd2), whose are rescued specifically by only the sno2&3 for Lsd1, and no snoRNA for Lsd2. This variability of rescue of different genes by the re-expression of the different snoRNA indicates rather a complex regulatory mechanisism controlled by the snoRNAs within the epithelium of the gut.

Considered altogether, in an overall holistic view, at the organismal level, these multiple results demonstrate a tight relationship between the expression of the snoRNA within the enterocytes controlling the expression of several genes, whose allow a precise regulation of several metabolic parameters. Moreover, these results suggest that the defects induced by the lack of the 3 snoRNAs within the gut could be compensated by the re-expression of them in the fat body, a result that reinforce the fact that the neurodegenative lesions as well as in parallel the longevity, are rather a consequence of the deregulation of the metabolic parameters, as TG and sterol. As the brain permanently needs a precise regulation of various nutrients to maintain its homeostasis all along the life, the deregulation of one of several of them, particularly chronically, as TG and sterol, leads to the development of age-progressive neurodegenerative lesions. In brief, we have shown that the brain lesions observed in aged flies are a consequence of the perturbation of the gut homeostasis, indicating a gut-brain axis.

## Materials & Methods

### Drosophila lines

*Drosophila melanogaster* flies were grown on standard medium (1.04% agar, 8% cornmeal, 8% brewer yeast and 2% nipagin as a mold inhibitor) at 25°C, 12:12 light:dark cycle in a humidity-controlled incubator. For ageing experiments, 15 adult female flies were crossed with 10 males per vial, and were transferred to new food vials every 2 days. Wild-Type *Canton-S* (CS) flies were used as control. The results described in this study were obtained from females. All genotypes were outcrossed to the wild-type CS (cantonization) at least 6 times to homogenize the genetic background. RU486 (Mifepristone) (*Sigma-Aldrich*, Cat. #M8046) to induce the Gene-Switch activity was dissolved in ethanol and mixed into the media when preparing food vials. RU486 doses used were 25 µg.mL^-1^ final concentration. Myo1A-Gal4 was kindly provided by B.A. Edgar (Heidelberg, Germany), CG8997-GS from H. Tricoire (Paris, France). Mex-GS (in VK02 site), UAS-jou (inserted randomly on the third chromosome), UAS-sno2 (in VK27 site), UAS-sno3 (in VK27 site), UAS-sno2&3 (in VK27 site), and UAS-3snoRNAs (in attP2 site), have been generated in our laboratory (Soulé et al., 2020). These various transgenic lines were then introduced into the F4-deletion genetic background by standard genetic crosses.

### Lifespan analysis

Following amplification, flies were harvested every day after hatching. Female flies were maintained with males in fresh food vials for 4 days at a density of 25 individuals per vial. On day 4, males were removed, while females were placed in a cage (about 200 to 300 females per cage), as in Soulé et al. (2020). The ageing animals were transferred to fresh food every two days, and the number of dead flies was scored. The lifespan plots were generated by calculating the percentage of survivorship every two days and plotting viability as a function of time (days) using log-rank test (Yang, et al. 2011).

### Quantitative RT-qPCR

Expression level of each snoRNA was measured on dissected gut or whole flies from 7-days-old female flies, as in Soulé et al. (2020). Quantitative RT-qPCR was performed on a QuantStudio-3 instrument (Applied Biosystem/ThermoFisher). For the snoRNA, we use a Taqman probe for each snoRNA (ThermoFisher). All assays were done in triplicate. Data were analyzed according to the ΔΔCt method, and normalized to RP49 levels. For the encoding genes, primers used are summarized in the Supplementary Table S1.

### Brain histology

#### a) Paraffin section

Flies were filled in collars and fixed for 4h in fresh Carnoy’s fixative (ethanol: chloroform: acetic acid at 30:15:5) as in (Soulé & Martin, 2020). Quantification of neurodegenerative lesions was performed on the entire brains using ImageJ software. Data are expressed as number of vacuolar/lesions per µm^2^ of measured surface.

#### b) Nile Red and BODIPY C11-581/591 labelling

All histological experiments were carried out on pre-fixed cryostat sections of the head from adult females aged of 7 days. For the Nile-Red, a stock solution at 10% (Sigma-Aldrich, Cat. #72485) is diluted in DMSO. After two washes of PBS-Tween 0.05% (PBST), the head sections are incubated for 30 min in the dark, with the Nile Red solution diluted 1/10 from an intermediate solution at 0,05% freshly prepared in PBST. The head sections are washed 2X with PBST, and mounted on glass slide in Mowiol. The brain are observed with a fluorescent microscope (Leica DM 600B) at two excitations wavelength of 499 nm and 543 nm, and at an emission of 519 and 600-650 nm, respectively. The photos are acquired with a digital camera (Hamamatsu C10600 ORCA-R2).

For the BODIPY C11-581/591 labelling, in contrast to other histological staining, the head of the flies are not fixed. The head sections are performed directly on frozen head, washed 2X with PBS, and incubated with the BODIPY C11-581/591 solution (D3861, ThermoFisher) diluted at 1/500 from a stock solution at 2µM, during 30 min. The head sections are washed 2X with PBST, mounted on glass slide in Mowiol, and observed in microscope using two different excitation wavelength, at 499 nm and 543 nm, and an emission at 519 et 600-650 nm, respectively. To calculate the ration of peroxidized versus non-peroxidized lipids, Image J (FIJI) is used. The intensity of the red and the green fluorescence is measured on the same image, and the ratio IntDen-Green/IntDen-Red is calculated for each measurement.

### Triglycerides measurement

The Triglycerides (TG) quantification has been performed with a colorimetric method using the enzymatic kit “LiquiColor TG kit” (Stanbio, Cat. #2200-430), as described in Soulé & Martin (2020). The TG determination is performed on 24 individual 7 day-old female fly.

### Sterol measurement

Sterols (including the cholesterol) in flies were measured as described in Soulé and Martin (2020). Measurements were carried out on 24 single 7-days-old adult female flies. Free sterol and sterol-ester levels were measured using the Amplex Red cholesterol assay kit (*Invitrogen*, Cat. #A12216), according to the manufacturer’s instructions. Notice that this enzymatic based-assay (kit) also measures all other sterols, and not solely the cholesterol (Serrano et al., 2024).

### *In-situ* hybridization (ISH)

For whole flies ISH, we used 5 day-old females, exactly as described in Soulé et al., (2020).

### Analysis of expression pattern of P[Gal4] and P[Gene-Switch] lines

All histological experiments were carried out on dissected guts from adult females aged of 7 days. To determine the expression pattern of Gal4 lines, samples were fixed in 4% PFA for 15 min, washed three times with PBS, and mounted in Mowiol. Images were collected using a Leica DM 600B light microscope (*Leica*, Germany), equipped with a Hamamatsu C10600 ORCA-R^2^ digital camera.

### Lipidomic Analysis

Lipid analysis was performed on the Functional Lipidomics Platform acknowledged by IBiSA (Infrastructure in Biology, Health and Agronomy) at ENSA, Lyon, exactly as described in (Jaque-Cabrera et al., 2025).

### Statistical analyses

Statistical comparisons were done with GraphPad Prism. Data were analyzed using the student-T test. Significance levels in figures were represented as *p < 0.05, **p < 0.01, ***p < 0.001. All quantitative data are reported as the mean ± S.E.M. (Standard Error of the Mean). Lifespan assays were subjected to survival analysis (log-rank test) using the freely available OASIS software (Yang et al., 2011). Supplementary Table S2 for the detailed statistics.

## Acknowledgments

We thank L. Mellottée for her technical assistance. H. Tricoire (Paris, France), and B.A. Edgar (Heidelberg, Germany) for fly stocks. This work was supported by the ANR (Agence Nationale de la Recherche, Ageing-jou, France), and by the CNRS (France).

## Author Contributions

JRM conceived and designed the experiments. SAI, TG, PD, and NBH performed experiments and analyzed data. JRM wrote the manuscript with input from all authors.

## Conflicts of Interest

The authors declare no conflict of interests.

## Supplementary Information

Ten Supplementary Figures and two Supplementary Tables are available online.

## Data Availability

The authors declare that all the data and the methods used in this study are available within this article. Supplementary Information are available from the corresponding authors upon reasonable request.

## Notes

### Competing Interest Statement

The authors have declared no competing interest.

