## Supplementary Figures and Tables for "A cluster of three snoRNAs including jouvence required in the gut determines lifespan and confers neuroprotection through metabolic parameters"

### Supplementary Figures & Tables

Suppl. Figure 1)

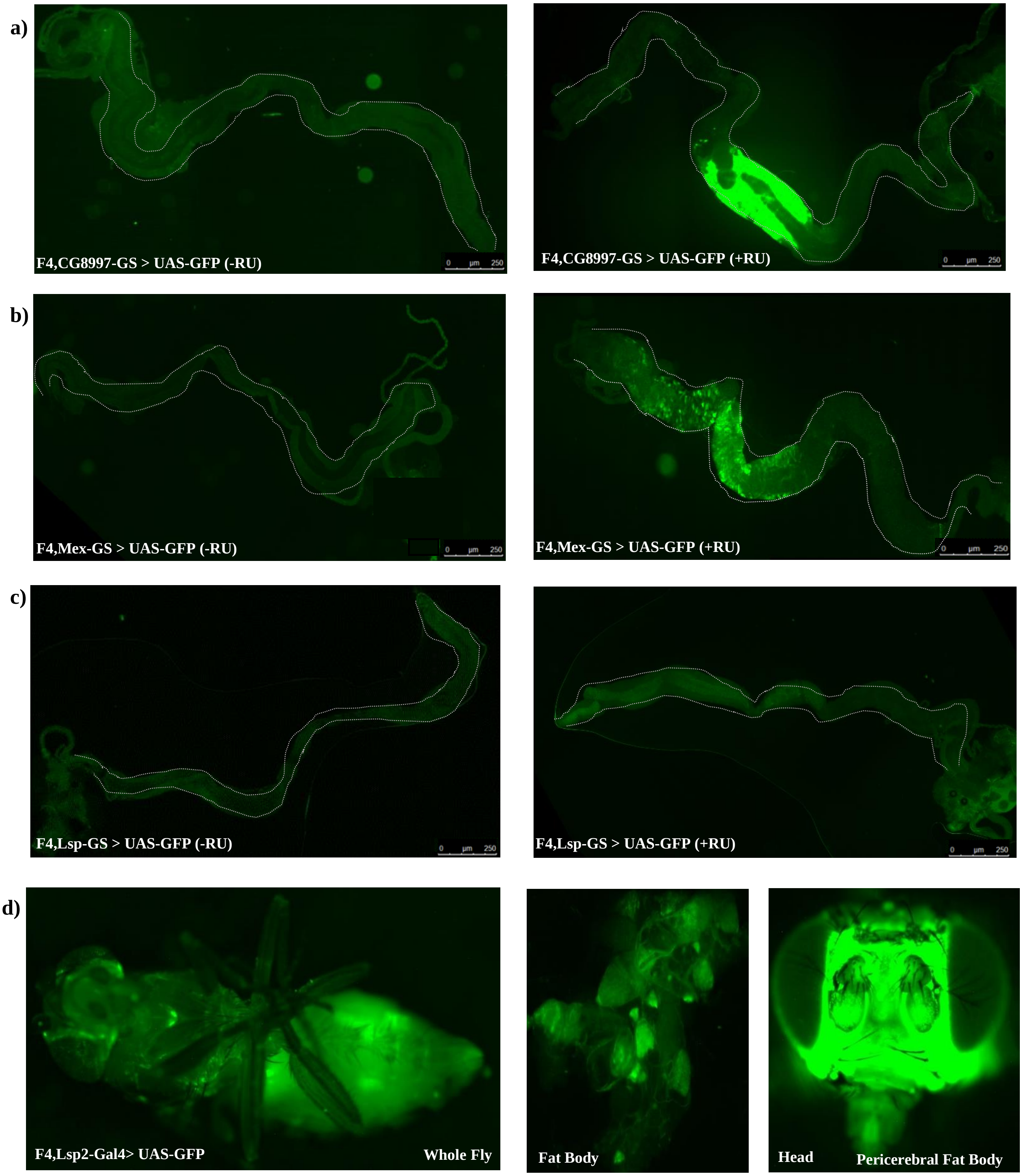

**Suppl. Figure 1) Expression Pattern of drivers CG8997-GS, Mex-GS, Lsp-GS and Lsp2-Gal4, in F4-Deleted flies.**

**a)** CG8997-GS shows GFP expression in enterocytes in the presence of RU486, predominantly in the R3 and R4 regions. **b)** Mex-GS drives GFP expression in enterocytes upon RU486 induction, with a possible expression in other cell types. **c)** No GFP expression in the gut with Lsp-GS. **d)** Lsp2-Gal4 drives GFP expression in the pericerebral and abdominal fat body.

**Suppl. Figure 2)**

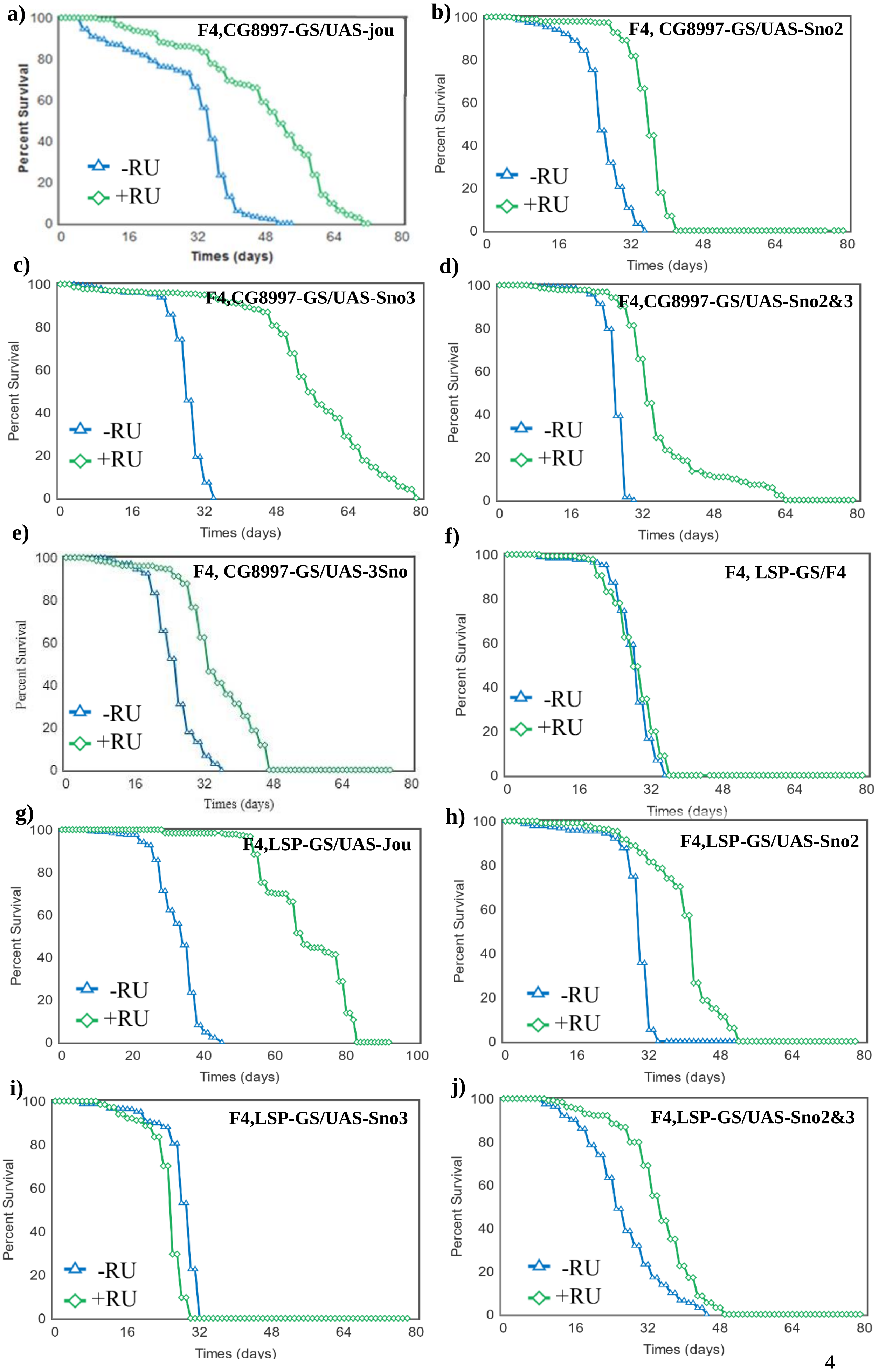

**Suppl. Figure 2) Targeted expression of each snoRNA in the enterocytes or in the fat body increases lifespan.**

Longevity test results (survival curve - decreasing cumulative) of the targeted expression of each snoRNA specifically in the enterocytes or in the fat body in F4-deleted flies compared to the F4-deleted flies whose don’t express any snoRNA. **a,b,c,d,e)** CG8997-GS driving each UAS-snoRNA or UAS-3snoRNAs in enterocytes fed with and without RU486, is sufficient to increase lifespan, with a stronger effect observed with the sno3 (c). **f)** F4, Lsp-GS/F4, fed with and without RU486, showing that the RU don’t have any effect on longevity. **g,h,i,j)** Lsp-GS driving each UAS-snoRNA in the Fat Body fed with and without RU486, is sufficient to increase lifespan, with a negative effect observed with the sno3 (i). For the number of flies, age in days at % mortality, and detailed statistics, see Supplementary Table-S2. P-value calculated by log-rank test using OASIS Sofware (Yang et al., 2011).

**Suppl. Figure 3)**

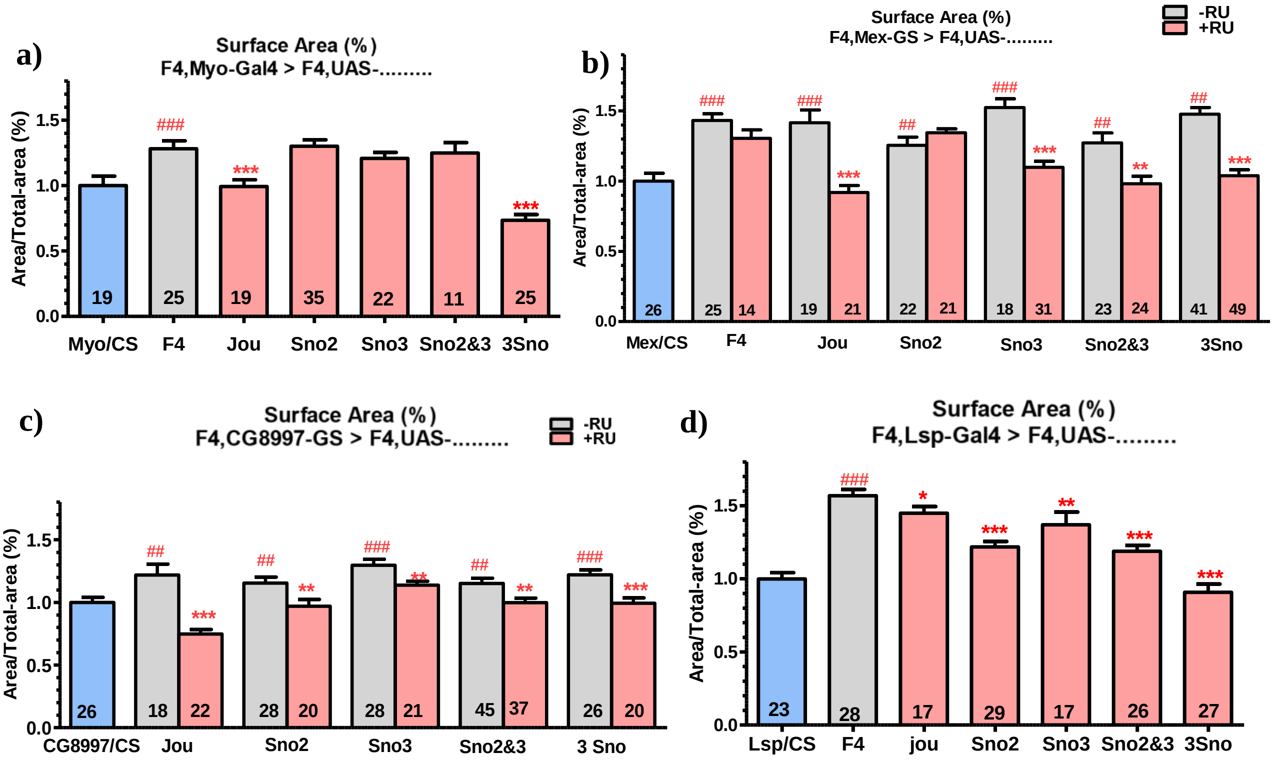


**Suppl. Figure 3) Quantification of Surface Area of neurodegenerative lesions in old flies.**

Each expression (putative rescue) is performed in the F4-deleted genetic background in old (40 day-old) flies. **a)** With the Myo1A-Gal4 driver, the re-expression of jou and/or the 3snoRNAs is sufficient to decrease the surface Area with a stronger effect observed with the 3 snoRNAs. **b)** The surface Area for the Mex-GS driving each UAS-snoRNA in the enterocytes, without and with feeding RU486 to induce the expression of the UAS-snoRNA transgenes. Re-expression of each snoRNA in enterocytes is sufficient to decrease the surface area of lesions with no effect observed with the sno2, while feeding the flies with RU, without snoRNA don't have any effect. **c)** Similar, but using the driver line CG8997-GS. The expression of sno2 using this driver is able to reduce the surface Area. **d)** Similar, but using the driver line Lsp2-Gal4 to target the snoRNA in the fat body. Again here, the stronger effect is observed with the 3 snoRNAs. Numbers within the histograms indicate the number of flies, errors bars represent the mean ± S.E.M, *p<0.05 ; **p<0.005 ; ***p<0.0005). p-value were calculated using the student-T test (Prism). Asterisks indicate significant differences compared to flies that never received RU (gray), while hashtags (#) highlight significant differences compared to wild-type flies (Blue) (Myo/CS, Mex/CS, CG8997-GS/CS, and Lsp/CS).

**Suppl. Figure 4)**

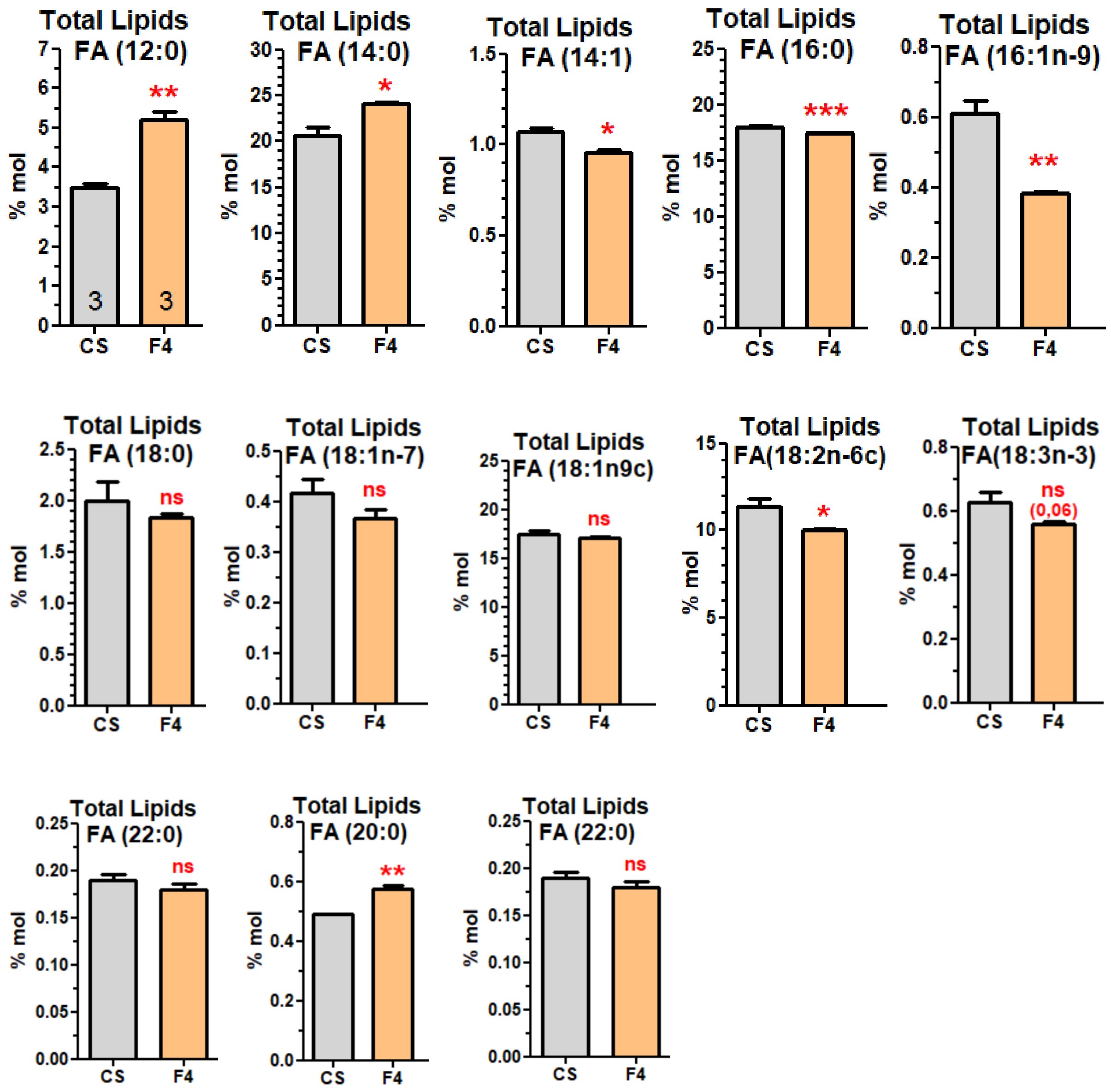


**Suppl. Figure 4) Fatty-Acid (FA) composition of Total Lipids.** The short FA (12:0 and 14:0) except (20:0), are increased, while several long FA (>16:0) are decreased.

**Suppl. Figure 5)**

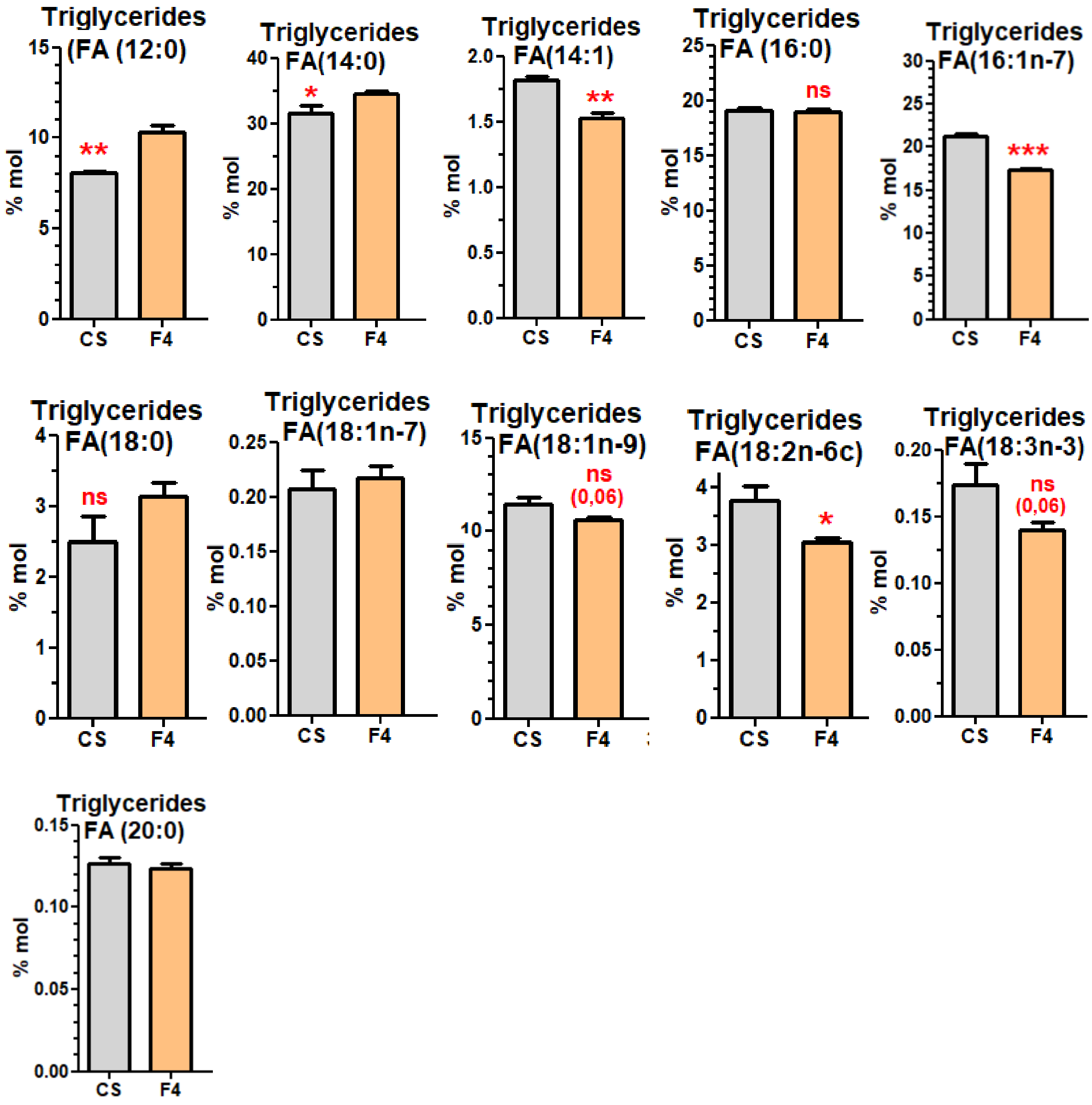


**Suppl. Figure 5) Fatty-Acid (FA) composition of Triglycerides.** The short saturated FA (12:0 and 14:0) are increased, while some monounsaturated or long FA (14:1, 16:1n-7, and >16:0) are decreased.

**Suppl. Figure 6)**

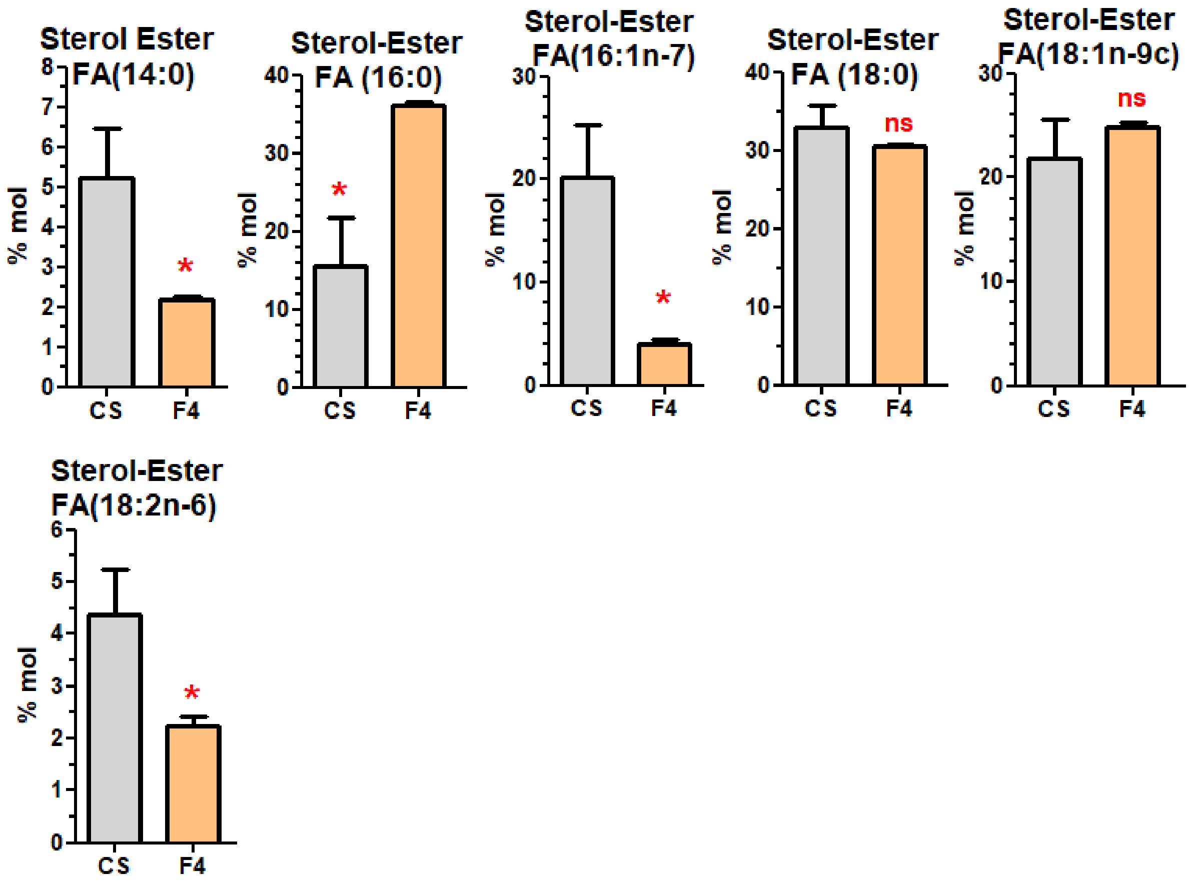


**Suppl. Figure 6) Fatty-Acid (FA) composition of Ester of Sterols.** Several FA are decreased, while only the 16:0 is increased.

Suppl. Figure 7)

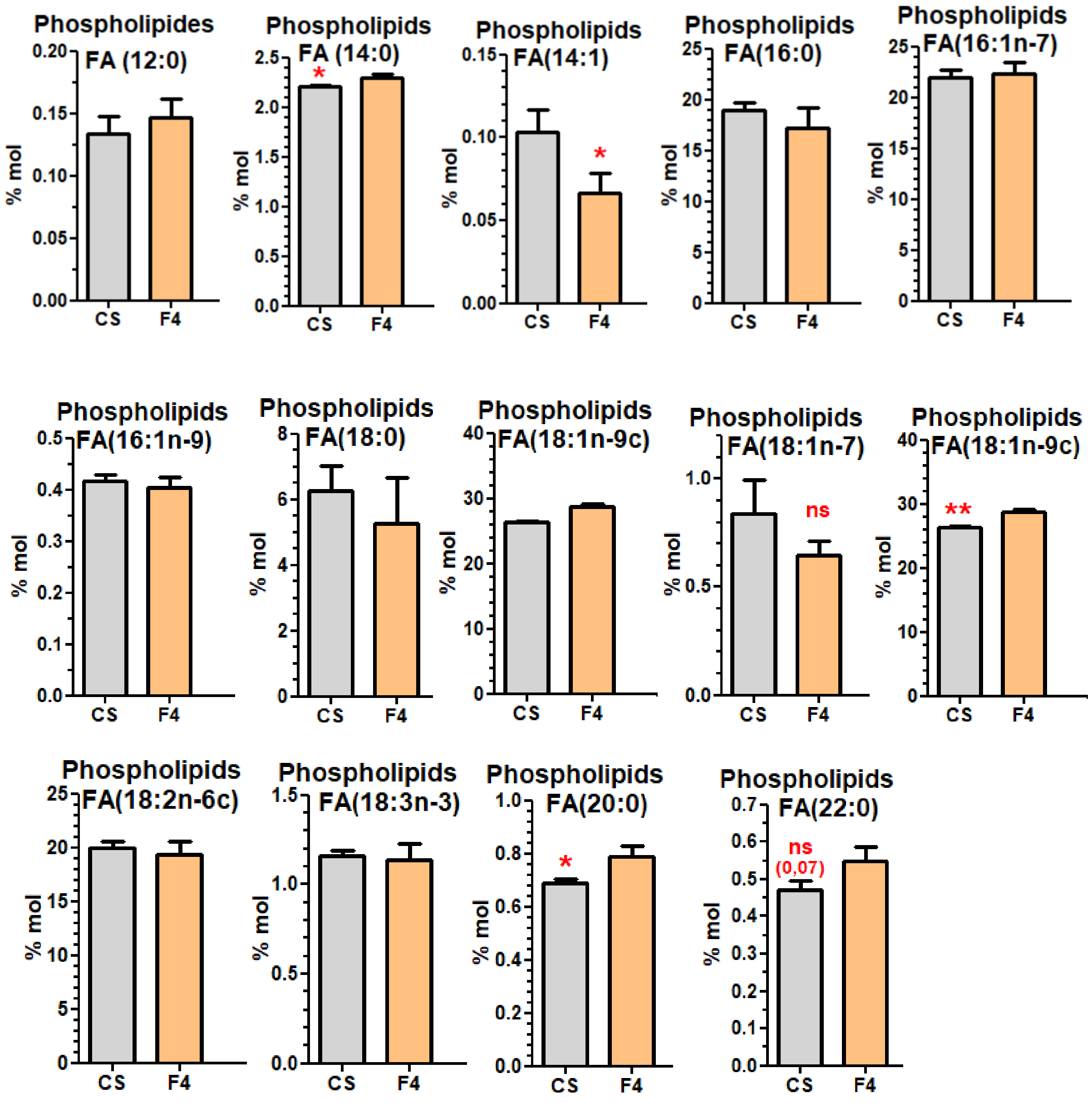


**Suppl. Figure 7) Fatty-Acid (FA) composition of Phospholipids.** The majority of FA are not modified.

**Suppl. Figure 8) Old flies (30 day-old)**

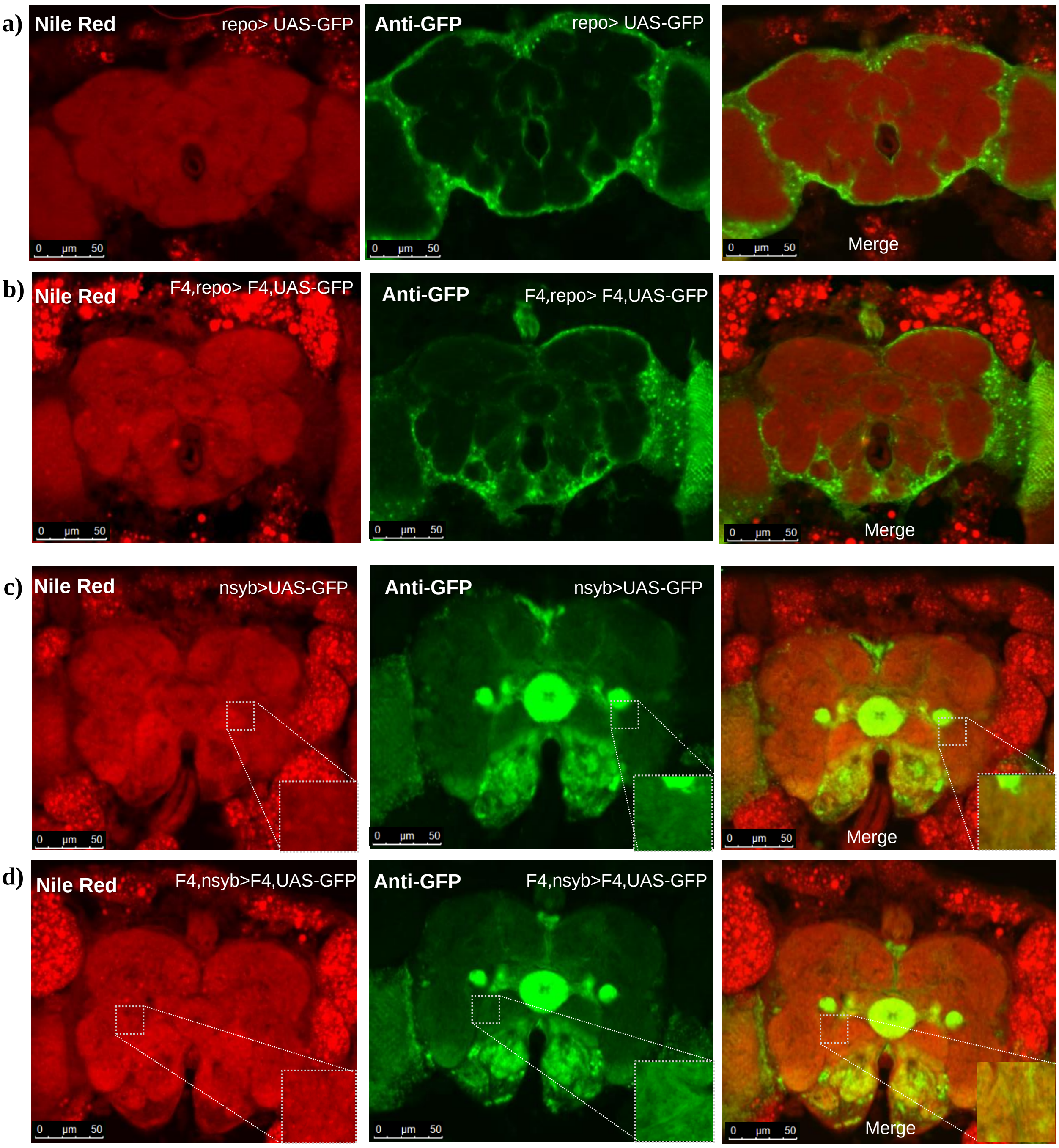


**Suppl. Figure 8) The Nile-Red overlaps with the neuronal tissue but not with the glial cells in old flies.**

**a,b,c,d)** First column, Nile Red Staining. Second column, Labelling of the glial cells using repo-Gal4>UAS-GFP or the neuronal cells using the n-syb-Gal4>UAS-GFP. Third column, Overlay.

#### Suppl. Figure 9)

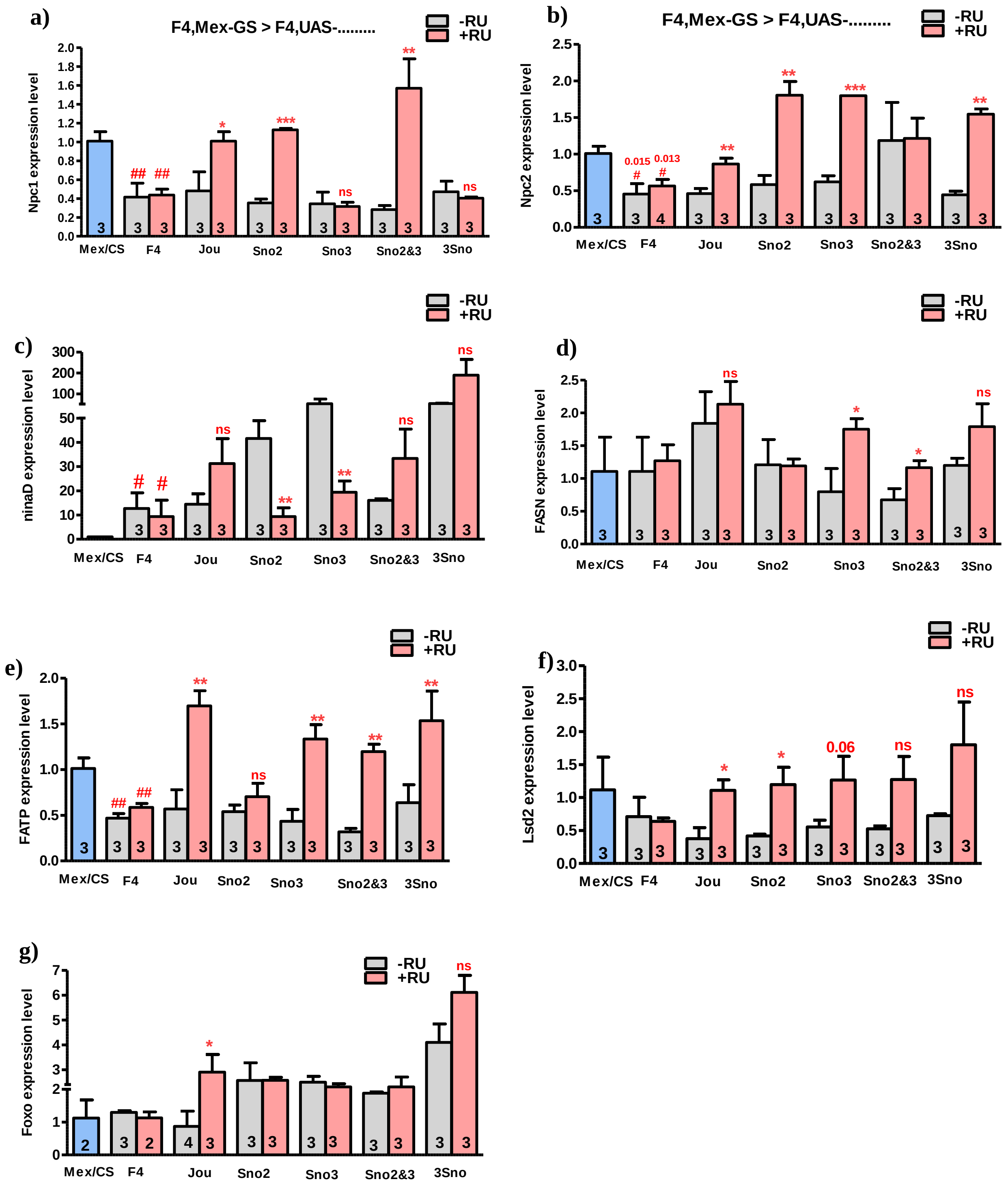


**Suppl. Figure 9) Different encoding genes are differentially rescued by the snoRNA driven by Mex-GS (RT-qPCR).**

RT-qPCR (SybGreen) of 7 different encoding genes performed on dissected gut of Controls, F4-deleted flies and targeted expression of each snoRNA in the enterocytes driven by Mex-GS.

**a)** NPC1 gene is decreased in F4, and is rescued by jou, sno2 and sno2&3, but not by sno3 neither by the 3 snoRNAs. **b)** NPC2 is decreased in F4, and is rescued by jou, sno2, sno3, sno2&3, and by the 3 snoRNAs **c)** ninaD is increased in F4, and is rescued by sno2, sno3. **d)** FASN is not decreased in F4, and its expression is increased by sno2, sno3 and the 3 snoRNAs. **e)** FATP is decreased in F4, and is rescued by the jou, sno2, sno3, sno2&3, and the 3 snoRNAs. **f)** Lsd2 (perilipin-2) is decreased in F4, but it is rescued by jou, sno2, sno3, sno2&3, and the 3 snoRNAs. **g)** FOXO is not decreased in F4, and it is increased only by jou, and the 3 snoRNAs. Statistics: The expression level of each gene in Control Wild-Type (Mex/CS) is used as reference (fixed to 1). Three independent biological replicates were done (n=3). (p-values) (* p<0,05; ** p<0,005; *** p<0,0005). Errors bars represent the mean +/- S.E.M. (p-value were calculated using the student T test, using Prism).

**Suppl. Figure 10)**

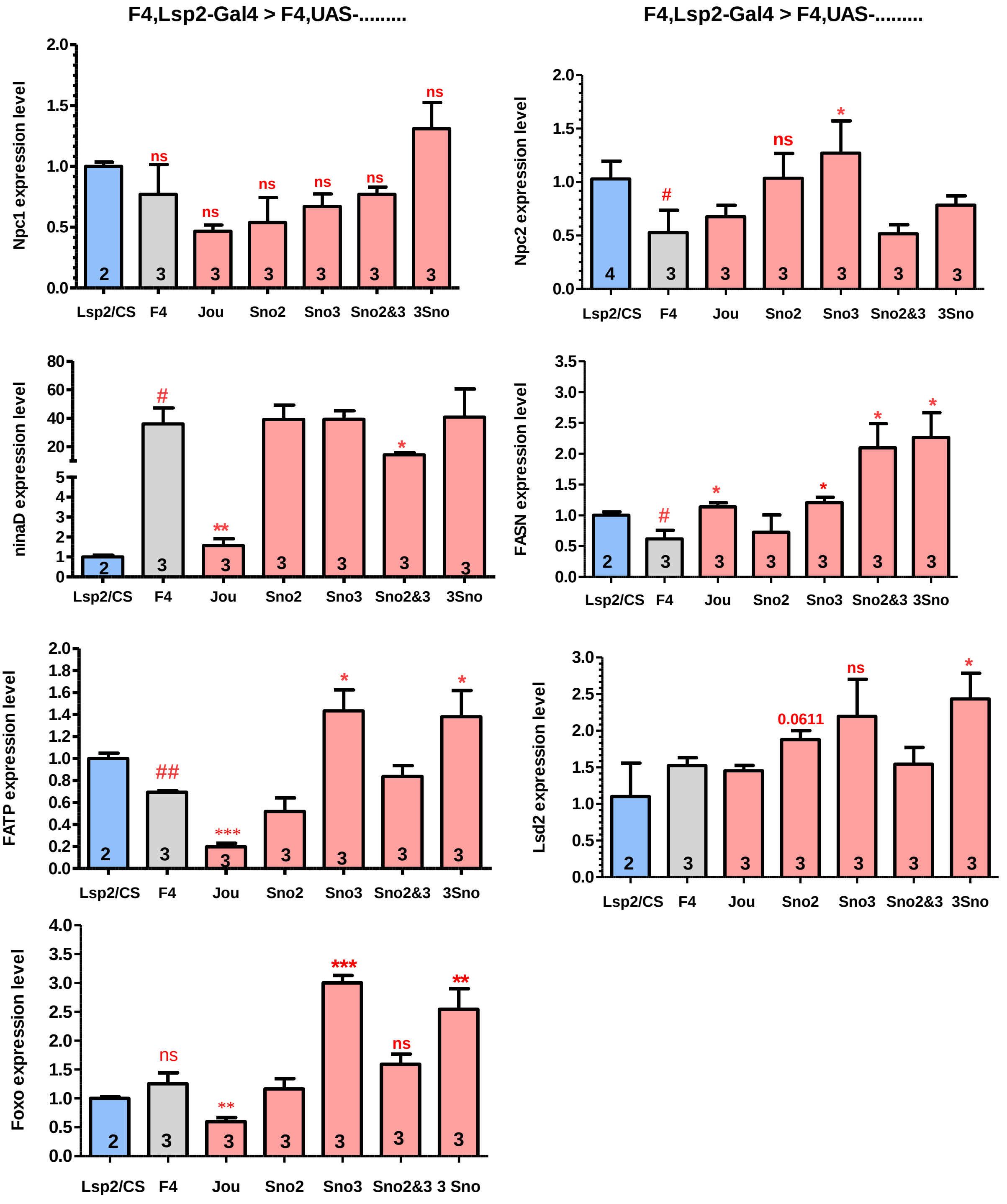


**Suppl. Figure 10) Different encoding genes are differentially rescued by the snoRNA driven by Lsp-Gal4 (RT-qPCR).**

RT-qPCR (SybGreen) of 7 different encoding genes performed on Whole flies of Controls, F4-deleted flies and targeted expression of each snoRNA in the enterocytes driven by Lsp2-Gal4.

| Gene | Forward Primers 5’-3’ | Reverse Primers 5’-3’ | Source |
|---|---|---|---|
| **RP49** | -AAGGGTATCGACAACAG- | -TTACTCGTTCTCCTTGAGA- | - |
| **FAS** | -CGTACGACCCCTCTGTTGAT- | -AGTGCAAGTTACCGGGAATG- | |
| **FATP** | -TACATCTACACCTCCGGCAC- | -GGGCGTGTAGAAGATGTCCT- | - |
| **Foxo** | -AGGCGCAGCCGAATAGACGAATTTA- | -TGCTGTTGACCAGGTTCGTGTTGA- | |
| **Lsd1** | -GTCAAGCTCTTCGAGCCATC- | -CACGAACTTATCGCCGACTG- | - |
| **Lsd2** | -AACGGAACAACTGGCAATGG- | -GGTCCAGTTTGGTGACGAAG- | - |
| **ninaD** | -CCAACAAAAGGCCTGGGTCT- | -CCGCCCCACATTTACCTTC- | - |
| **NPC1a** | -CCTATTTCGTCATCATGGCAGC- | -CCAGTATAACAACCAGGGATGC- | - |
| **NPC2a** | -GTACGCGGTAATTGCCTGTG- | -TACGACTCGTCCTTCTCCAAC- | - |

**Suppl. Table S1) Sequences of primers used for SYBR Green qPCR**. The “Gene” column lists official gene names according to FlyBase. The “Forward Primer” and “Reverse Primer” columns show primer sequences for each gene, while the “Source” column indicates their origin. If no reference is provided, primers were designed in the laboratory. The highlighted primers in the first row (Rp49) correspond to the reference gene.

#### Suppl. Table S2) Longevity Statistics: number of flies, age in days at % mortality, and statistics.

| Genotypes | | Age in days at % mortality | | | | | Statistics | |
|---|---|---|---|---|---|---|---|---|
| Lines | Number | 25% | 50% | 75% | 90% | 100% | Genotypes | p-value |
| **F4, Myo/F4** | **256** | **29** | **31** | **35** | **37** | **43** | | |
| **F4,Myo/UAS-jou** | **335** | **37** | **43** | **47** | **55** | **63** | **F4,Myo vs jou-8M** | **0,000** |
| **F4, Myo/UAS-sno-2** | **216** | **43** | **47** | **51** | **55** | **55** | **F4,Myo vs sno2** | **0,000** |
| **F4,Myo/UAS-sno-3** | **235** | **40** | **44** | **46** | **50** | **52** | **F4,Myo vs sno3** | **0,000** |
| **F4, Myo/UAS-sno-2,3** | **237** | **40** | **42** | **48** | **52** | **56** | **F4,Myo vs sno2&3** | **0,000** |
| **F4,Myo-Gal4/ UAS-3 Sno** | **149** | **37** | **39** | **45** | **51** | **53** | **F4,Myo vs 3 Sno** | **0,000** |
| **F4,Mex/F4 (-RU)** | **243** | **25** | **27** | **27** | **29** | **29** | **F4,Mex/F4 (-RU) VS F4,Mex/F4 (-RU)** | **0,270** |
| **F4,Mex/F4 (+RU)** | **220** | **20** | **26** | **30** | **32** | **32** | | |
| **F4,Mex-GS/UAS-jou (-RU)** | **252** | **30** | **32** | **34** | **36** | **42** | **F4,Mex/jou (-RU) VS F4,Mex/jou (+RU)** | **0,000** |
| **F4,Mex-GS/UAS-jou (+RU)** | **236** | **35** | **39** | **39** | **43** | **48** | | |
| **F4,Mex-GS/UAS-sno-2 (-RU)** | **270** | **46** | **46** | **48** | **50** | **58** | **F4,Mex/sno2 (-RU) VS F4,Mex/sno2 (+RU)** | **0,00** |
| **F4,Mex-GS/UAS-sno-2 (+RU)** | **213** | **51** | **63** | **67** | **73** | **77** | | |
| **F4,Mex-GS/UAS-sno3 (-RU)** | **195** | **40** | **42** | **44** | **46** | **50** | **F4,Mex/sno3 (-RU) VS F4,Mex/sno3 (+RU)** | **0,00** |
| **F4,Mex-GS/UAS-sno3 (+RU)** | **203** | **51** | **65** | **73** | **81** | **85** | | |
| **F4,Mex-GS/UAS-sno2,3 (-RU)** | **252** | **26** | **28** | **30** | **32** | **38** | **F4,Mex/sno2&3 (-RU) VS F4,Mex/sno2&3 (+RU)** | **0,00** |
| **F4,Mex-GS/UAS-sno2,3 (+RU)** | **255** | **31** | **35** | **37** | **50** | **62** | | |
| **F4,Mex-GS/UAS-3 Sno (-RU)** | **202** | **22** | **26** | **28** | **32** | **32** | **F4,Mex/3 Sno (-RU) VS F4,Mex/3 Sno (+RU)** | **0,00** |
| **F4,Mex-GS/UAS-3 Sno (+RU** | **109** | **26** | **28** | **34** | **38** | **38** | | |
| **F4,Lsp-Gal4/ F4** | **299** | **15** | **17** | **19** | **23** | **25** | | |
| **F4,Lsp-Gal4/UAS-jou** | **300** | **19** | **23** | **25** | **29** | **31** | **F4,Lsp vs jou-8M** | **0,00** |
| **F4,Lsp-Gal4/UAS-sno2** | **248** | **26** | **28** | **28** | **30** | **32** | **F4,Lsp vs sno2** | **0,00** |
| **F4,Lsp-Gal4/UAS-sno3** | **178** | **29** | **31** | **33** | **37** | **39** | **F4,Lsp vs sno3** | **0,00** |
| **F4,Lsp-Gal4/UAS-sno2&3** | **111** | **21** | **25** | **27** | **29** | **33** | **F4,Lsp vs sno2&3** | **0,00** |
| **F4,Lsp-Gal4/3 Sno** | **169** | **19** | **25** | **27** | **29** | **31** | **F4,Lsp vs 3 Sno** | **0,00** |

#### Suppl. Table S2 - continued

| | | | | | | | | |
|---|---|---|---|---|---|---|---|---|
| F4,CG8997-UAS,jou (-RU) | 219 | 27 | 35 | 37 | 41 | 54 | F4,CG8997-GS/jou (-RU) VS F4,CG8997-GS/jou (+RU) | 0,00 |
| F4,CG8997-UAS,jou (+RU) | 144 | 39 | 51 | 59 | 63 | 71 | | |
| F4,CG8997-UAS,sno2 (-RU) | 284 | 23 | 25 | 29 | 33 | 35 | F4,CG8997-GS/sno2 (-RU) VS F4,CG8997-GS/sno2 (+RU) | 0,00 |
| F4,CG8997-UAS,sno2 (+RU) | 191 | 34 | 36 | 38 | 40 | 42 | | |
| F4,CG8997-GS/UAS-sno3 (-RU) | 224 | 26 | 28 | 30 | 32 | 34 | F4,CG8997-GS/sno3 (-RU) VS F4,CG8997-GS/sno3 (+RU) | 0,00 |
| F4,CG8997-GS/UAS-sno3 (+RU) | 222 | 51 | 55 | 65 | 73 | 79 | | |
| F4,CG8997-GS/UAS-sno2,3 (-RU) | 318 | 26 | 26 | 28 | 28 | 30 | F4,CG8997-GS/sno2&3 (-RU) VS F4,CG8997-GS/sno2&3 (+RU) | 0,00 |
| F4,CG8997-GS/UAS-sno2,3 (+RU) | 224 | 31 | 33 | 37 | 52 | 64 | | |
| F4,CG8997-GS/UAS-3 Sno (-RU) | 187 | 22 | 26 | 28 | 32 | 36 | F4,CG8997-GS/3 Sno (-RU) VS F4,CG8997-GS/3 Sno (+RU) | 0,00 |
| F4,CG8997-GS/UAS-3 Sno (+RU) | 205 | 31 | 33 | 43 | 47 | 47 | | |
| F4,Lsp-GS/F4 (-RU) | 164 | 25 | 29 | 31 | 33 | 35 | F4,Lsp-GS/F4 (-RU) VS F4,Lsp-GS/F4 (+RU) | 0,55 |
| F4,Lsp-GS/F4 (+RU) | 136 | 26 | 28 | 32 | 34 | 36 | | |
| F4,Lsp-GS/UAS-jou (-RU) | 327 | 28 | 34 | 36 | 38 | 45 | F4,Lsp-GS/jou (-RU) VS F4,Lsp-GS/jou (+RU) | 0,00 |
| F4,Lsp-GS/UAS-jou (+RU) | 189 | 58 | 68 | 80 | 83 | 83 | | |
| F4,Lsp-GS/UAS-sno2 (-RU) | 211 | 28 | 30 | 32 | 32 | 34 | F4,Lsp-GS/sno2 (-RU) VS F4,Lsp-GS/sno2 (+RU) | 0,00 |
| F4,Lsp-GS/UAS-sno2 (+RU) | 215 | 36 | 42 | 44 | 50 | 52 | | |
| F4,Lsp-Gs/UAS-sno-3 (-RU) | 266 | 28 | 30 | 30 | 32 | 32 | F4,Lsp-GS/sno3 (-RU) VS F4,Lsp-GS/sno3 (+RU) | 0,00 |
| F4,Lsp-Gs/UAS-sno-3 (+RU) | 315 | 24 | 26 | 28 | 28 | 30 | | |
| F4,Lsp-GS/UAS-sno2,3 (-RU) | 264 | 21 | 25 | 31 | 37 | 45 | F4,Lsp-GS/sno2&3 (-RU) VS F4,Lsp-GS/sno2&3 (+RU) | 0,00 |
| F4,Lsp-GS/UAS-sno2,3 (+RU) | 129 | 31 | 35 | 39 | 43 | 49 | | |
